# Citric acid water effects on mouse health, motivation, and performance in virtual reality locomotion-based tasks

**DOI:** 10.64898/2026.08.04.742856

**Authors:** Brenna E. Manuel, Grayson O. Sipe

## Abstract

Traditionally, complex behavioral tasks in mice have relied upon water restriction as an external motivator to increase task engagement. However, citric acid (CA) water, the technique whereby water is given *ad libitum* but made sour by the addition of CA, has emerged as an alternative to typical water restriction. Evidence suggests CA water can effectively motivate task performance while improving animal welfare in alignment with the 3Rs of animal research and reducing experimenter labor. While promising, the applicability of CA water in mice remains incompletely characterized with higher concentrations only tested in rats and “ramping” schedules, where mice progress to increasingly higher concentrations, indirectly examined. Here, we evaluate four CA concentration/schedule combinations for their effects on mouse health (weight changes, home cage behaviors, fecal counts) and motivation to drink regular water (lick counts, drinking behaviors) in female and male C57BL/6J mice. We find that a schedule ramping from 1% to 2% CA after one week is the easiest for mice to adapt to and sustained 2% CA use maintains robust lick counts for at least 5 weeks. Additionally, CA has been directly characterized for wheel-turning and touchscreen tasks, but not virtual reality (VR) tasks, an increasingly popular class of behavioral experiments. Therefore, we also assess how 2% CA affects motivation and task performance in two VR treadmill tasks (running and stopping task). We find that CA water does not improve task performance above that of mice given regular water, but does limit competing motivations and produce more uniform, reward-motivated behavior.

## Introduction

Complex decision-making tasks in rodents produce rich behavioral readouts that enable experimenters to probe higher-level cognitive processes such as instrumental learning and spatial navigation (Juavinett et al., 2018). However, rodents typically require external motivation to encourage them to learn complex task criteria. Water restriction is one of the most common motivation methods in which home cage water access is limited to motivate animals to correctly perform behavioral tasks through earning water rewards (Goltstein et al., 2018; Guo et al., 2014; Rowland, 2007). While water restriction is effective at increasing task learning and performance, this method requires high experimenter diligence to avoid ethical health concerns, as weight must be carefully monitored and supplemental water given to maintain physiological health, even during days when no behavior is performed (e.g. weekends). In more extreme schedules, water restriction can result in animal distress and/or unintended abnormal physiology that confounds experimental interpretation (Barkus et al., 2022; Willems, 2009). To reduce experimenter labor and permit rodents to self-regulate water intake, researchers are investigating alternatives to traditional water restriction in accordance with the Refinement principle of the 3Rs framework for animal research.

Citric acid (CA) water is one alternative external motivation paradigm in which home cage water is always accessible but made sour by the addition of CA, the common food preservative found naturally in citrus fruits (Reinagel, 2018; Urai et al., 2021; Watson et al., 1986). CA reduces water palatability and rodents will self-restrict their consumption. While first characterized in rats, recent studies have demonstrated that CA water use in mice produces motivation comparable to water restriction for a wheel-turning task without causing adverse health effects (Urai et al., 2021). Additionally, CA water can be flexibly implemented through different concentrations and access schedules (e.g., weekends only, ramping from 1% to 2% CA, etc). Consequently, CA water as a motivator has expanded to a variety of tasks and experimental setups (Guillaumin et al., 2023; Ishizu et al., 2024; Kondo et al., 2025). Despite progressively widespread use of CA water, different laboratories have used different CA concentrations and access schedules, and the consequence of these differences on task performance and motivation have not been characterized. For example, 2% CA is the most commonly reported percentage used in mice, but other researchers have documented using between 0.4% and 3% CA (Dong et al., 2025; Gershon et al., 2025; Mai et al., 2024). Researchers implemented these concentrations either by maintaining a single percentage or by successively “ramping” to higher percentages. Additionally, 4% CA water has been used frequently in rats (Bao et al., 2023; Reinagel 2018, Watson et al., 1986), but to our knowledge, has not been assessed for use in mice. Therefore, it is crucial to thoroughly assess the relative health and motivation effects for different commonly reported CA parameters. Here, we characterize the longitudinal motivation and health effects of 1% - 4% CA water, expanding upon the foundational work by Urai et al. (2021) by including higher CA percentages and extending the long-term characterization of 2% CA. We also assess two ramping schedules of different rates of increase to determine whether gradually increasing CA percentages over time produces increased motivation, as ramping schedules have been used to maintain motivation throughout an experiment (Mai et al., 2024).

Finally, while CA-based motivation has been characterized for touchscreen tasks (Dzinic et al., 2026) and wheel-turning tasks (Urai et al., 2021), the effectiveness of CA water for motivating locomotion-based tasks has not been directly assessed. Nonetheless, researchers already use CA water as a motivator for both head-fixed and freely moving locomotion tasks (Dong et al., 2025; Jordan et al., 2021; Rupprecht et al., 2024), including virtual reality (VR) treadmill navigation tasks (Heiser et al., 2025). As closed-loop VR treadmill tasks become increasingly popular and accessible (Thurley & Ayaz, 2017), with many paradigms continuing to rely on traditional water restriction (Harvey et al., 2009; Henschke et al., 2020; Pakan et al., 2018; Radvansky & Dombeck, 2018), establishing the effectiveness of CA water in these tasks would thus expand the potential repertoire of behavioral paradigms in which CA water can confidently be applied. Accordingly, we assess the motivational influences of CA water on two VR treadmill tasks: one where mice are trained to run toward increasingly farther reward patches and one where mice are trained to stop in reward patches. Altogether, our study provides a more comprehensive understanding of both the applicability and limitations of CA water allowing for other experimenters to make more-informed decisions on their behavioral methodologies.

## Results

### The effects of CA percentages and schedules on weight change and home cage behaviors

To examine how citric acid (CA) concentration and schedule affect the health and motivation of mice, we conducted a series of group-housed experiments where we exchanged the home cage drinking water of mice for various concentrations of CA water (Fig. 1 and Table 1). The 0% CA cohort received regular water while the 2% CA cohort received 2% CA water. The slow ramp (SR) and fast ramp (FR) cohorts were successively transitioned to higher CA percentages across time: 1% increase every week for the SR and 1% increase every 3 days for the FR. We also utilized a smaller 4% CA cohort to test the effects of starting mice directly on 4% CA water. We checked the health and weight of each mouse in each cohort daily and removed mice that had lost more than 20% of their starting weight from the study along with their group-housed cage mates.

**Figure 1.**
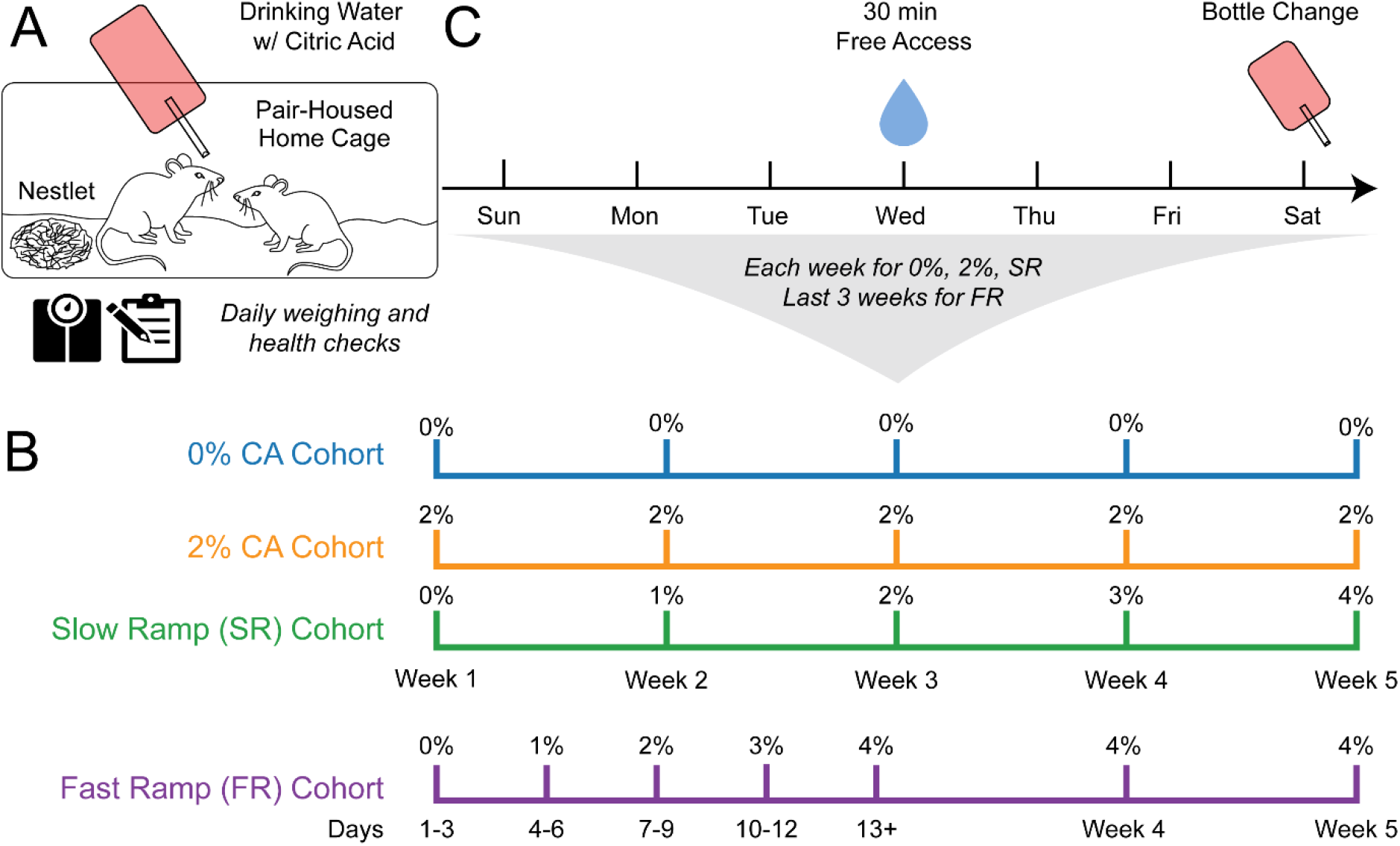
Experimental design and timelines for the 5-week characterization cohorts. *A*, Schematic of pair-housed home cage environment and measurements. Regular cage water was replaced with citric acid water of cohort-specific concentrations. *B*, For 5 weeks, 4 cohorts on different CA percentages and schedules were weighed and checked daily. Timelines indicate CA concentrations for each cohort across the 5 weeks. 0% CA is equivalent to regular water. *C,* On Wednesdays, mice were separated for 30 minutes and given unrestricted access to regular water. This occurred every week for the 0% CA, 2% CA, and SR cohorts and every week starting in week 3 for the FR cohort. Bottles were changed every Saturday except for the FR cohort which had bottles changed every 3 days.

**Table 1.**
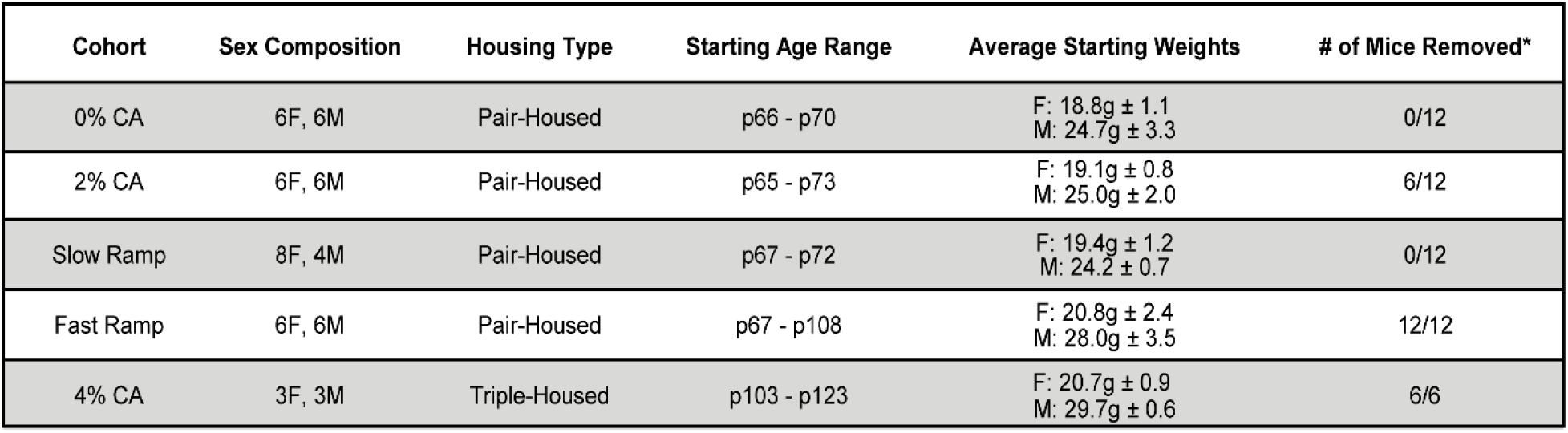
Composition of the group-housed characterization cohorts. A total of 54 mice (29F, 25M) were used for 5 different CA water schedules (0% CA being a control cohort). Aside from the triple-housed 4% cohort, mice were all pair-housed. The starting ages and weights are from day 0 (0%, 2%, 4% CA) or day 1 (SR, FR) of each experiment. *More mice were removed from the study than reached the 20% weight loss threshold due to group-housed mice having to be removed from the study together.

Consistent with the normal weight gain of 2–3-month-old C57BL/6J mice (The Jackson Laboratory, 2015), the 0% CA mice steadily gained weight above their baseline at a rate of 1.3% ± 0.1% a week (Fig. 2*A*). Three of the 2% CA mice crossed the 20% weight loss threshold and were removed from the study along with their pair-housed cage mates (6/12 mice removed) (Fig. 2*B*). The 6 remaining mice (4F, 2M) gained weight across the 5 weeks at a rate comparable to the controls (1.4% ± 0.3%) (Fig. 2*F*). Conversely, the SR cohort gradually lost weight across weeks (-3.0% ± 0.3%) but did not have any mice drop below the weight loss cutoff (Fig. 2*C*, *F*). Moreover, this weight loss diminished over time, as indicated by the average rate of weight loss being better represented by an exponential decay than a linear decline (Extended Data Fig. 2-1*A*). The FR mice also continually lost weight, but while on 2% CA water, 4 mice dropped below the weight loss threshold (8/12 mice removed), so the experiment was terminated on day 9 (Fig, 2*D*). Similarly, all 6 mice in the small 4% CA cohort exceeded 20% weight loss within 3 days, and the experiment was terminated (Fig. 2*E*).

**Figure 2.**
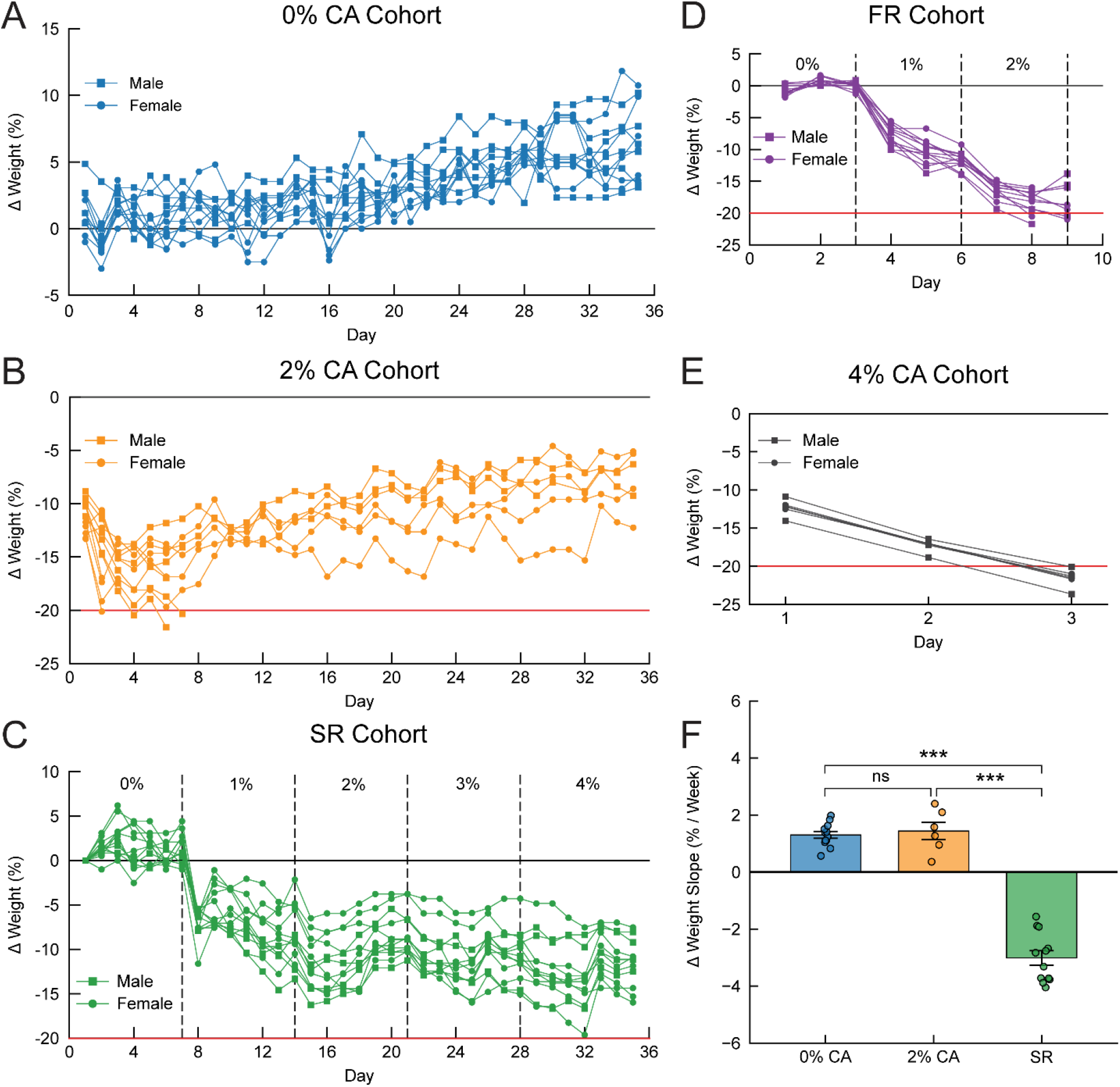
Cumulative body weight changes across different CA concentrations and schedules*. **A,*** Total weight change across time normalized to baseline weight for the 0% CA cohort. ***B-E,*** Depict the same as ***A*** but for the 2% CA, SR, FR, and 4% CA cohorts and normalized to their baseline weights respectively. Red lines at -20% denote the weight loss threshold for removing mice from the study. ***F,*** Total weight-change rates derived from individual linear regression slopes and compared across cohorts. Kruskal-Wallis test, (*H*(2) = 20.98, *p* = 2.78 × 10^−5^). Post hoc Dunn’s tests, 0% vs 2% CA (*p* = 0.78, CI: [−0.82, 0.51]); 0% vs SR (*p* = 1.49 × 10^−4^, CI: [3.69, 4.98]); and 2% CA vs SR (*p* = 6.44 × 10^−4^, CI: [3.68, 5.30]). For more information see Extended Data 2-1.

We next quantified daily weight fluctuations to obtain weight change measures with higher temporal resolution. Longitudinal tracking revealed mice in the 2% CA cohort initially lost weight for the first 3-4 days before they gained weight and stabilized (Fig. 3*A*). This suggests the first 4 days are an important habituation period for 2% CA. Because weight gain on day 5 was potentially influenced from the prior day’s regular water access (Extended Data Fig. 2-1*C*), the subsequent weight recovery trajectory of 2% CA alone cannot be distinguished. We then examined the daily weight fluctuations of the SR cohort and found that each CA concentration increase was followed by a sharp weight loss the next day (Fig. 3*A*). However, the magnitude of these losses successively decreased with increasing CA concentrations such that the decrease following the 3% to 4% CA transition was not significantly different from that following the 2% to 3% CA transition (Extended Data Fig. 2-1*B*). This data provides evidence of CA schedule habituation in the SR cohort that supports the previous findings of total weight change stabilization.

**Figure 3.**
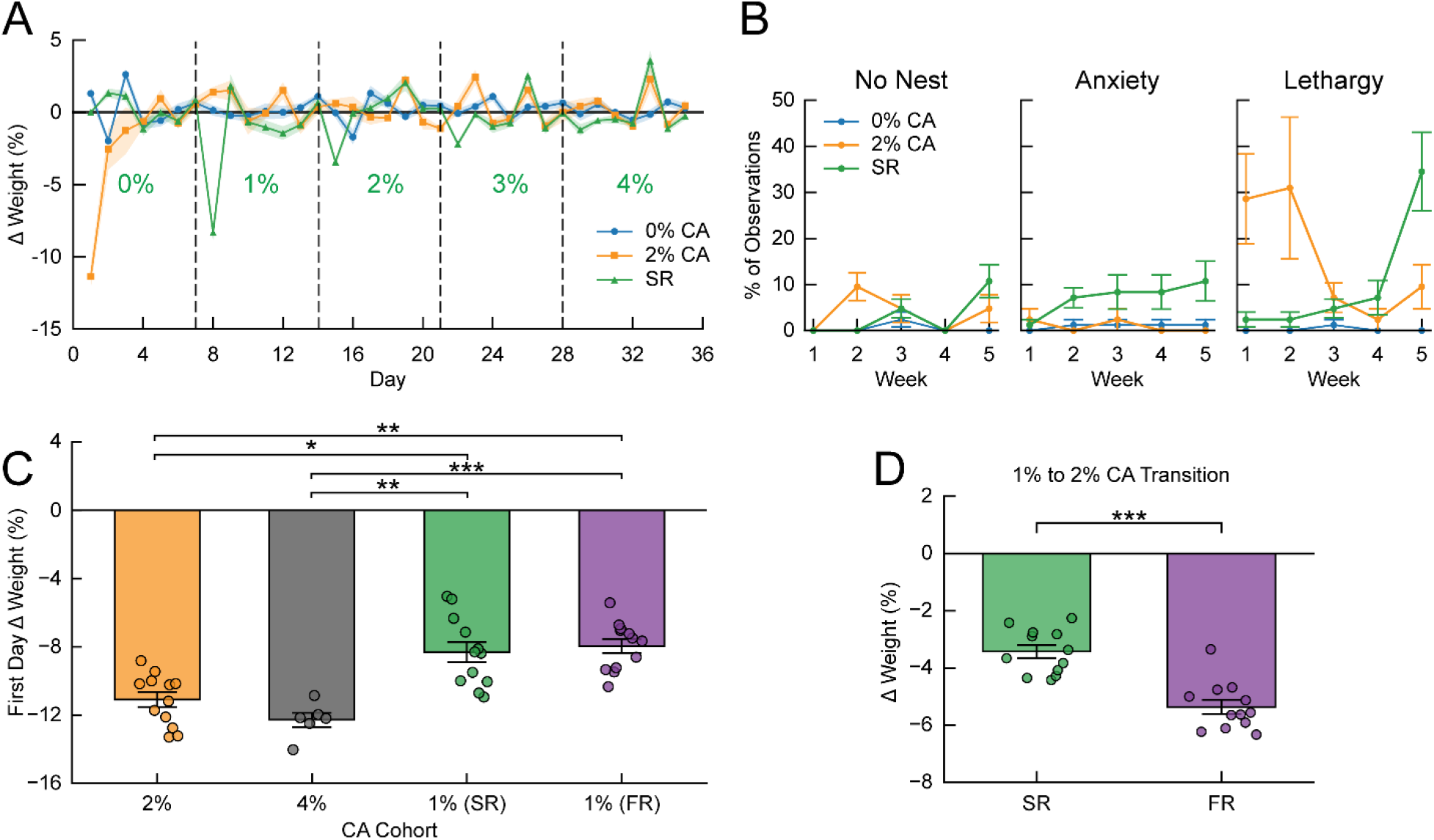
Comparisons of daily weight changes and home cage behaviors across CA concentrations and schedules. ***A*,** Daily weight changes for the 0% CA, 2% CA, and SR cohorts as a comparison of one day relative to the next. ***B,*** Relative frequency of aberrant home cage behaviors observed during daily weighing for the 0% CA, 2% CA, and SR cohorts. Nest-making F1-LD-F1 test, cohort effect (*F*(1.69, 18.95) = 7.51, *p* = 0.0055; week effect (*F*(1.80, ∞) = 4.68, *p* = 0.012); cohort-week interaction (*F*(2.19, ∞) = 2.94, *p* = 0.048). Mann-Whitney U post hoc test, 0% vs 2% CA (*p* = 0.0028, CI: [-5.71, -2.86]). Anxiety F1-LD-F1 test, cohort effect (*F*(1.77, 21.54) = 6.11, *p* = 0.0098); week effect (*F*(3.12, ∞) = 0.97, *p* = 0.41); cohort-week interaction (*F*(4.12, ∞) = 1.67, *p* = 0.15). Mann-Whitney U post hoc test, 0% CA vs SR (*p* = 0.015, CI: [-11.43, 0.00]. Lethargy F1-LD-F1 test, cohort effect (*F*(1.48, 9.59) = 7.26, *p* = 0.016); week effect (*F*(3.59, ∞) = 1.68, *p* = 0.16); and cohort-week interaction (*F*(4.52, ∞) = 3.30, *p* = 0.0075). Mann-Whitney post hoc tests, 0% vs 2% CA (*p* = 5.72×10^−4^, CI: [-27.14, -4.29]); 0% CA vs SR (*p* = 5.76×10^−4^, CI: [-11.43, -2.86]). ***C,*** Relative weight losses for the day after each cohort first started on CA water (transition from 0% CA). Kruskal-Wallis test, (*H*(3) = 23.66, *p* = 2.94 × 10^−5^). Significant post hoc Dunn’s tests, 2% CA vs FR (*p* = 0.0030, CI: [-4.46, -1.89]); 4% CA vs FR (*p* = 6.47 × 10^−4^, CI: [-5.35, -3.09]); 2% CA vs SR (*p* = 0.011, CI: [-4.25, -1.15]); and 4% CA vs SR (*p* = 0.0030, CI: [-5.66, -2.20]). ***D,*** A comparison of the relative weight losses after the 1% to 2% CA transition for the SR and FR cohorts. Mann-Whitney U test, (*p* = 1.96 × 10^−4^, CI: [1.29, 2.71]). For more information see Extended Data 2-1.

To compare the impact of starting CA concentration on initial weight loss severity, we next calculated the magnitude of initial weight decreases from 1%, 2%, and 4% CA (Fig. 3*C*). The average weight loss on the first day after starting CA water provides a measure of the initial palatability challenge posed by each concentration, independent of whether mice subsequently habituated to that CA concentration. The SR and FR mice, which both transitioned first to 1% CA, exhibited similar initial weight losses. Moreover, these decreases were smaller than those observed in both the 2% CA and 4% CA cohorts, whose initial weight decreases were also comparable despite 4% CA water being twice as concentrated. This may indicate a potential floor effect of initial weight loss. Because the SR and FR cohorts shared both an initial 0% to 1% CA transition and a subsequent 1% to 2% CA transition, we then isolated the effects of ramp rate from concentration increases by directly comparing this 1% to 2% CA transition. While both cohorts exhibited similar weight losses after the initial 0% to 1% CA transition (Fig. 3*C*), FR mice had significantly greater weight losses after the subsequent 1% to 2% CA transition (Fig. 3*D*). This finding indicates a dissociable effect of ramp rate from the CA concentration increases themselves, with the faster ramp rate leading to worse weight loss outcomes for subsequent CA transitions.

Because physiological weight metrics do not capture the full wellbeing of the mice, we also monitored qualitative home cage behaviors during daily weighing. For the 3 cohorts that continued for the full 5 weeks (0% CA, 2% CA, and SR), we observed relatively few aberrant home cage behaviors (Fig. 3*B*) and no additional signs of pain or distress such as hunching or complete immobility. However, when comparing the 2% CA and SR cohorts to the 0% CA mice, there were some notable behavioral differences. Nest building varied by week with only 2% CA mice exhibiting significantly delayed nest building. Contrastingly, only SR mice displayed significantly increased anxiety-like behaviors. For lethargy, 2% CA mice initially showed high lethargy (∼30% of observations) that decreased over time, while SR mice initially showed low lethargy that gradually increased over time, particularly with a sharp increase (from 7% to 34% of observations) during week 5 (4% CA). While derived from qualitative observations, these three metrics together detail more subtle indicators of schedule-specific distress phenotypes that are not apparent from weight data alone.

### Weekly free-water access reveals thirst-motivated licking behaviors

The 0% CA, 2% CA, and SR cohorts received 30 minutes of access to regular water once weekly to assess motivation to lick for regular water (Fig. 1*C*). We temporarily separated mice into empty cages and measured their licking via a 24-cage lickometer (Fig. 4*A, B*). At the end of each 30-minute session, we also measured fecal counts. As expected, the 2% CA and SR mice overall had higher lick counts than the 0% CA mice (Fig. 4*C*). Additionally, the lick counts for the 2% CA cohort remained stable across the 5 weeks, whereas the SR cohort lick counts increased for the first 3 weeks after which 3% and 4% CA produced no significant lick count increases relative to the time-matched 2% CA cohort. To assess if the body weight changes of individual mice were related to their lick counts across sessions, we next performed repeated measures correlations (Fig. 4*D, F, H*). The relationship between lick count and total weight change had a significant negative correlation for the 0% CA and SR mice, but no significant correlation for the 2% CA mice. This is consistent with how the 2% CA mice gained weight across weeks but maintained the same approximate lick counts.

**Figure 4.**
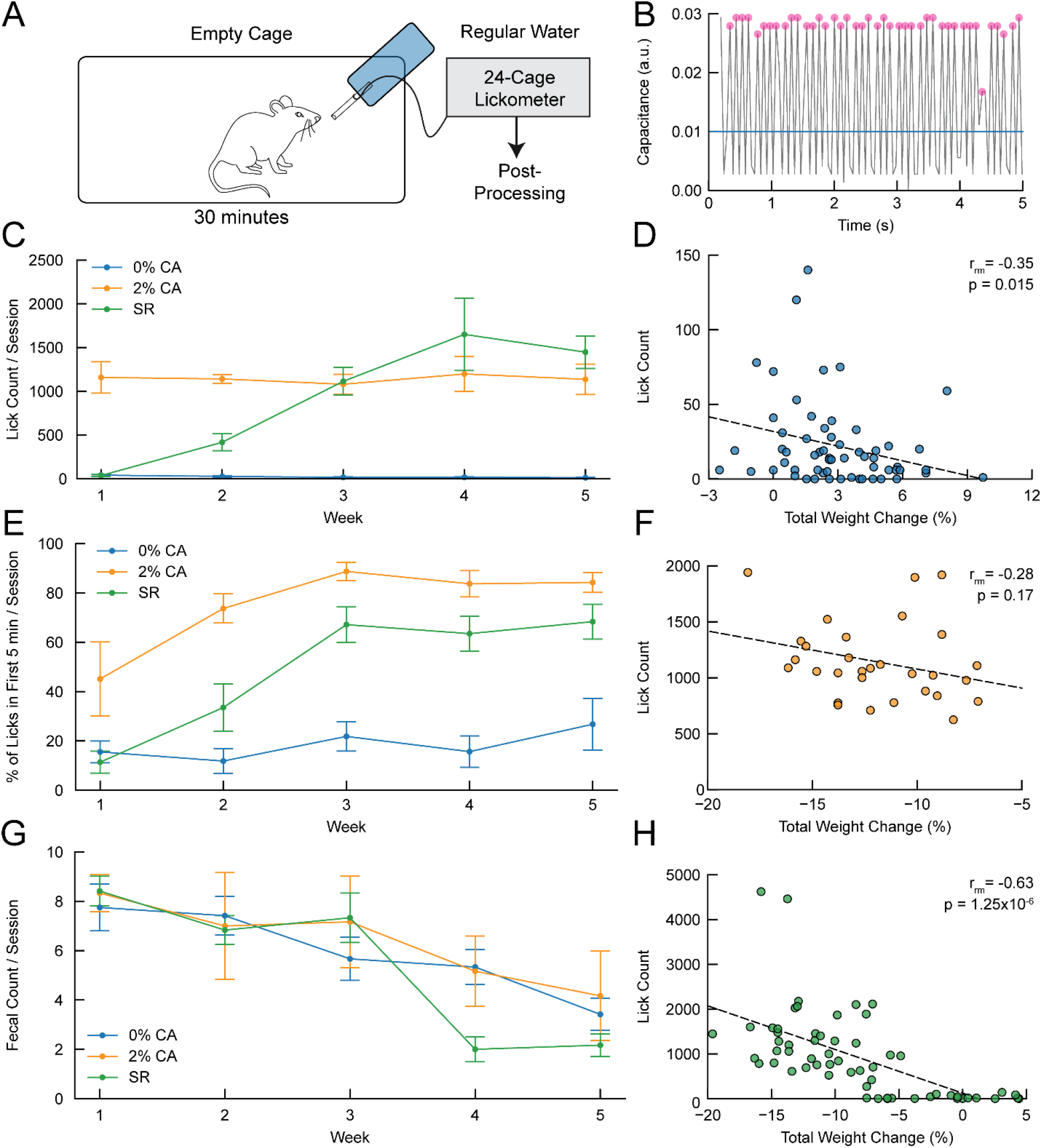
Thirst-motivated licking behaviors and fecal counts from weekly water access sessions. ***A***, Schematic of lick-detection setup used for the 5-week characterization cohorts. ***B,*** Sample post-processed data with the peak detection filter applied (2% CA, female, week 5). ***C***, Lick counts for the 0% CA, 2% CA, and SR cohorts during weekly 30-minute water-access sessions. F1-LD-F1 test, cohort effect (*F*(1.81, 25.56) = 55.01, *p* = 1.15 × 10^−9^); week effect (*F*(3.48, ∞) = 4.59, *p* = 0.0019); cohort-week interaction (*F*(5.38, ∞) = 14.11, *p* = 1.06 × 10^−14^). Mann-Whitney U post hoc tests for 2% CA vs SR, Week 3 (*p* = 0.96, CI: [-453.00, 344.00]); Week 4 (*p* = 0.96, CI: [-912.00, 614.00]); Week 5 (*p* = 0.15, CI: [-953.00, 318.00]). ***D,*** Repeated measures correlation between licks and baseline-normalized body weight changes for 0% CA mice (CI: [−0.57, −0.073], df = 47). ***E,*** Percentage of licks from ***C*** that occur within the first five minutes of the session, excluding sessions with fewer than 2 licks. F1-LD-F1 test, cohort effect (*F*(1.95, 15.91) = 27.37, *p* = 7.77 × 10^−6^); week effect (*F*(3.26, ∞) = 12.95, *p* = 5.43 × 10^−9^); and cohort-week interaction (*F*(5.42, ∞) = 2.98, *p* = 0.0088). Post hoc Mann-Whitney U tests for 2% CA vs SR, Week 3 (*p* = 0.034, CI: [1.57, 39.30]); Week 4 (*p* = 0.18, CI: [-3.15, 31.52]); Week 5 (*p* = 0.18, CI: [-5.15, 28.80]). ***F,*** Same as ***D*** but for 2% CA mice (CI: [-0.61, 0.13], df = 23). ***G,*** Fecal counts in cages after 30-minute lick sessions. F1-LD-F1 test, cohort effect (*F*(1.40, 8.32) = 0.32, *p* = 0.66); week effect (*F*(2.86, ∞) = 25.36, *p* = 2.40 × 10^−16^); and cohort-week interaction (*F*(4.11, ∞) = 2.69, *p* = 0.028). Post hoc Mann-Whitney U test for Week 4, 0% CA vs SR (*p* = 0.0040, CI: [1.00, 5.00]). ***H,*** Same as ***D*** and ***F*** but for SR mice (CI: [-0.77, -0.42], df = 47). For more information see Extended Data 2-1.

To further dissect drinking pattern differences, we next investigated temporal licking patterns for frontloading, a pattern of drinking in which rodents drink more of a rewarding substance at the start of access (Ardinger et al., 2022; Lardeux et al., 2013). Here, frontloading was defined as the percentage of overall licks occurring within the first 5 minutes of the session with a higher percentage indicating more frontloading (Fig. 4*E*). The 0% CA cohort had no clear frontloading pattern while the 2% CA and SR cohorts exhibited increased frontloading across time with 2% CA mice showing more frontloading than SR mice while both were on 2% CA water. There was no difference in frontloading between the 2% CA and SR mice for the last 2 weeks. As a simple supporting measure for both hydration and stress levels (Barkus et al., 2022; Calvo-Torrent et al., 1999; Fertig & Edmonds, 1969), we then analyzed weekly fecal count data from the water licking sessions. Fecal counts for the 0% CA, 2% CA, and SR mice decreased similarly across time, consistent with habituation effects (Fig. 4*G*). However, during week 4, SR mice had a significant drop in fecal count as compared to the 0% CA mice. This potentially indicates worsening dehydration that was not overtly apparent from the weight data.

### Mice on 2% CA do not perform better than mice on regular water in a VR running task

We designed two head-fixed virtual reality (VR) treadmill tasks to assess the motivational effects of CA water in locomotion-based tasks (Fig. 5) while further characterizing the weight changes of mice on 2% CA (Extended Data Fig. 5-1). The first was a running-focused task modeled after a progress ratio procedure, which is one of the most robust ways to measure motivation (Hodos, 1961; Johnson et al., 2022). This task exponentially increased the average distance between reward zones (RZs) every 15 rewards (i.e., 1 level progression). Thus, mice learned to increase their travel distances to keep earning rewards of 5% sucrose water (3µL drop). Six mice on regular water (0% CA) and 6 mice on 2% CA water engaged in self-paced training in the running task for 3 weeks (Table 2). While both groups of mice increased travel distances across weeks, there was a significant condition effect with 0% CA mice having a trend of traveling farther (Fig. 6*A*, Ext. Data Video 4). Correspondingly, 4 of 6 0% CA mice achieved higher level attainment than all 2% CA mice, which further indicates better task performance (Fig. 6*B*, Extended Data Fig. 6-1*A, B*). For additional performance metrics, we next analyzed reward counts across weekdays to assess within-week motivation fluctuations and across training weeks to assess changes over learning. We found a significant main effect of weekday and no significant condition main effect or condition-week interaction (Fig. 6*C*). However, post hoc comparisons revealed no significant differences between individual weekdays. Similarly, across training weeks, reward counts changed across time (main effect of week: *F*(1.45, ∞) = 5.07, *p* = 0.013) but did not differ between 0% and 2% CA mice (main effect of condition: *F*(1.00, 7.63) = 3.67, *p* = 0.093; condition-week interaction: *F*(1.45, ∞) = 1.23, *p* = 0.28). There were also no significant post hoc comparisons across training weeks. Together, these findings indicate that 0% and 2% CA mice did not differ in the number of rewards earned despite differences in running distance and level attainment. Finally, we examined post-reward licking and task acquisition rate as additional motivational metrics. We found that 2% CA mice more consistently licked the rewards that they received (Fig. 6*D*) and initially acquired the task at a rate comparable to 0% CA mice (Fig. 6*E*).

**Figure 5.**
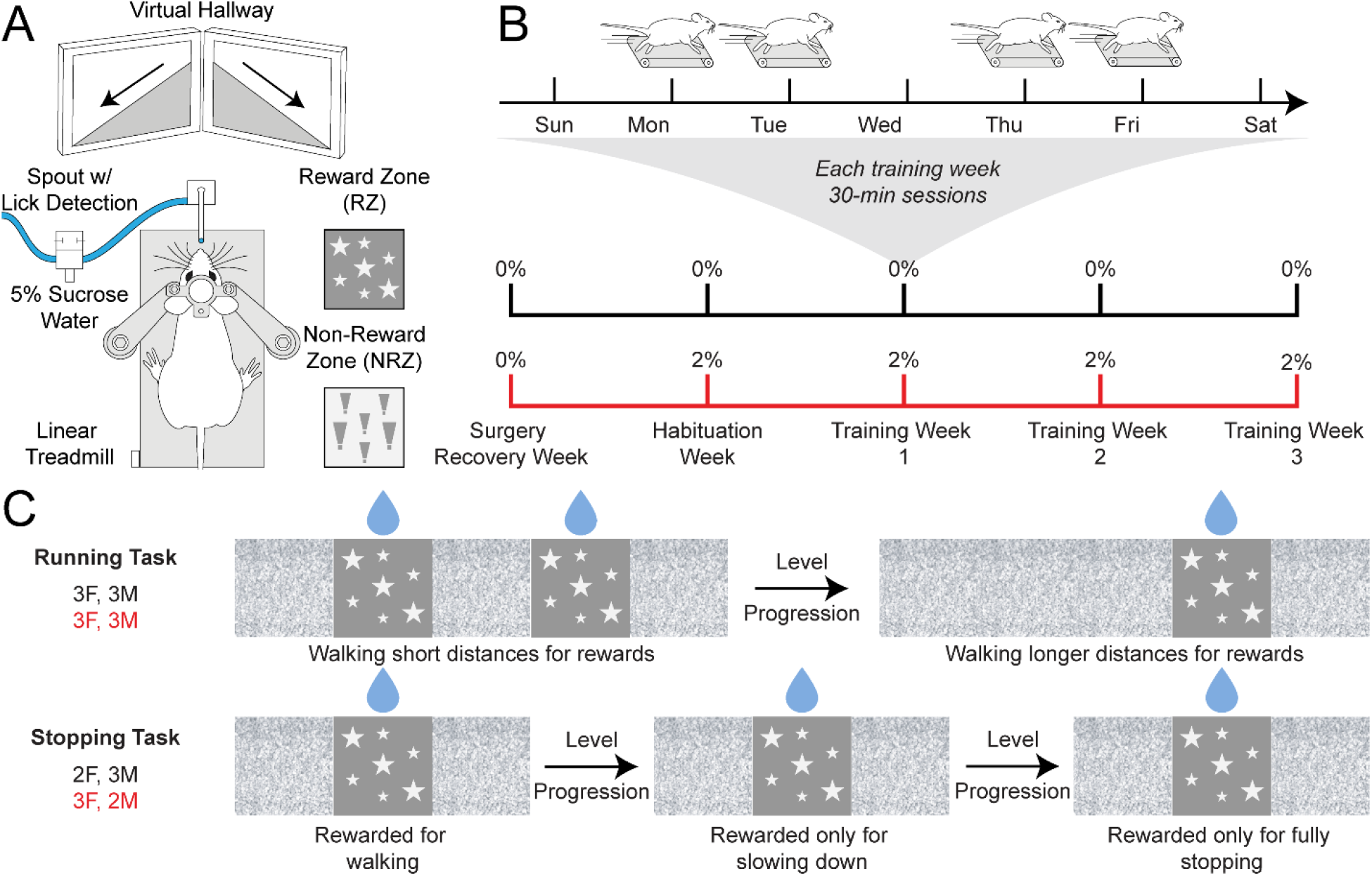
Virtual reality setup and timelines for the running and stopping tasks. ***A,*** Closed-loop virtual reality schematic used for both tasks and example wall textures. ***B***, General timelines for the VR cohorts. Mice first underwent headplate surgeries and were given a week to recover. The next week, half of each cohort started on 2% CA water while half remained on regular water. This week, termed habituation week, also consisted of experimental setup habituation. Mice then engaged in 3 weeks of self-paced training on one of 2 tasks. Training was 4 days a week, 30 minutes each day on the days indicated by mice on treadmills. ***C***, Simplified schematics of the two VR tasks. In the running task, mice initially traversed a virtual hallway where reward zones were spatially close together. Every level progression, the distance between reward zones increased exponentially. In the stopping task, mice were initially rewarded just for walking into reward zones. Then, mice were only rewarded for spending more time (i.e., slowing down) inside of reward zones. After further level progressions, mice were only rewarded for fully stopping inside of reward zones. All level progressions for both tasks were based on mouse reward counts, thus requiring mice to earn rewards to keep progressing. For weight change data from VR cohorts see Extended Data 5-1.

**Figure 6.**
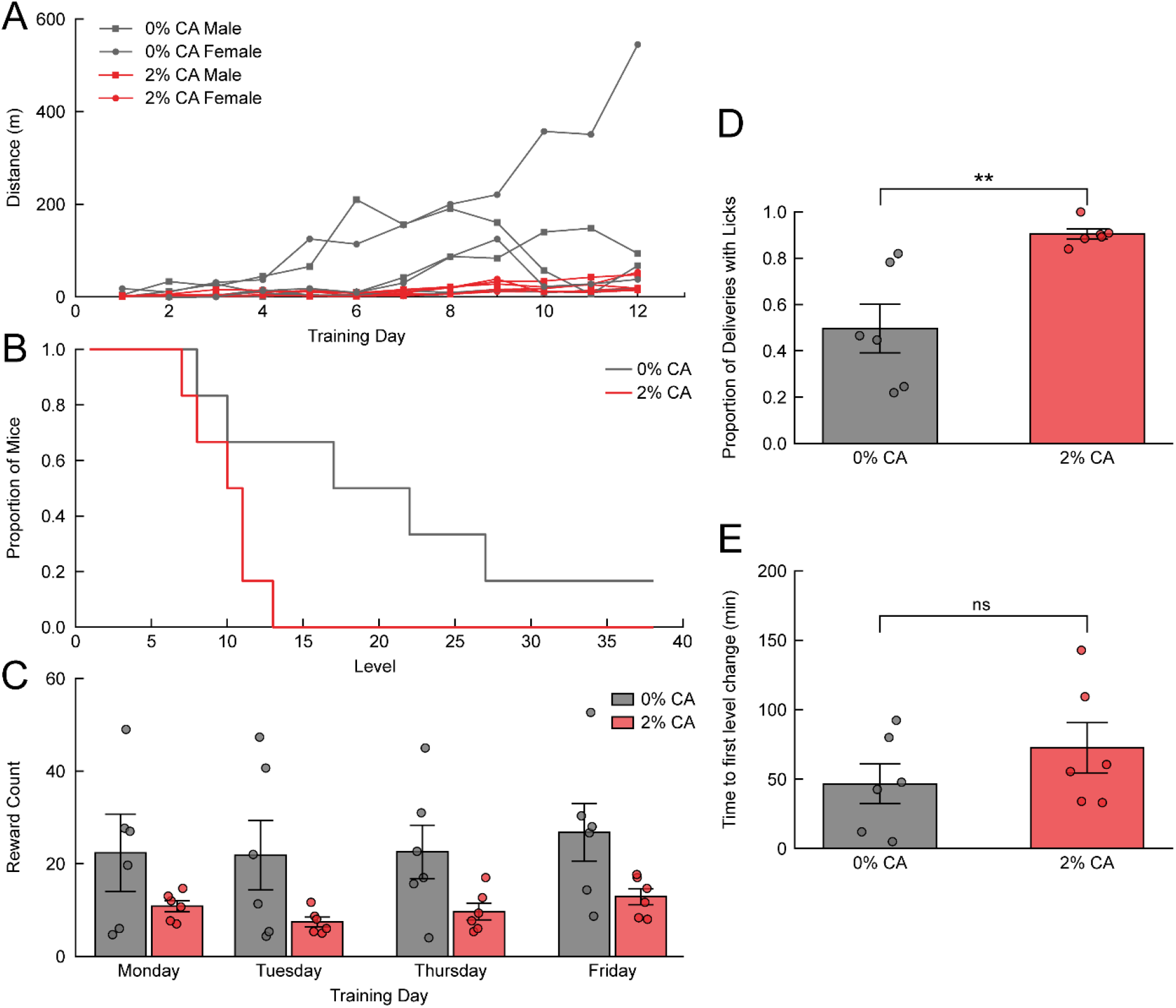
Across-session comparisons for the task performance and engagement results of the running task. ***A,*** Total distances traveled by individual 0% and 2% CA mice within the 30-minute training sessions of the running task. F1-LD-F1 test, condition effect (*F*(1.00, 8.62) = 5.28, *p* = 0.048); week effect (*F*(1.41, ∞) = 24.60, *p* = 1.01 × 10^−8^); and condition-week interaction (*F*(1.41, ∞) = 1.06, *p* = 0.33). Post hoc Mann-Whitney U, 0% vs 2% CA (*p* = 0.093, CI: [-2.98, 160.04]). ***B,*** A Kaplan-Meier-style plot depicting the level attainment of the 6 mice on 0% CA and 6 mice on 2% CA. Adjacent levels differed by 15 rewards. ***C,*** A breakdown by weekday and CA percentage for the average number of rewards attained on each training day. F1-LD-F1 test, condition effect (*F*(1.00, 7.20) = 3.41, *p* = 0.11); weekday effect (*F*(1.96, ∞) = 3.94, *p* = 0.020); condition-weekday interaction (*F*(1.96, ∞) = 0.32, *p* = 0.72). Post hoc Wilcoxon signed-rank tests, Mon. vs Tue. (*p* = 0.73, CI: [-3.83, 7.83]); Mon. vs Thu. (*p* = 0.73, CI: [-4.83, 4.83]); Mon. vs Fri. (*p* = 0.064, CI: [-5.50, -1.00]); Tue. vs Thu. (*p* = 0.31, CI: [-6.17, 1.17]); Tue. vs Fri. (*p* = 0.27, CI: [-11.00, 0.00]); and Thu. vs Fri. (*p* = 0.31, CI: [-7.67, 0.33]). ***D,*** Comparison collapsed across sessions of the proportion of reward deliveries that have at least one lick within 2 seconds of delivery. 0% vs 2% CA Mann-Whitney U test, (*p* = 0.0022, CI: [-0.66, -0.10]). ***E,*** Cumulative time across sessions for mice to progress from level 1 to level 2 as a proxy measure for initial task acquisition. 0% vs 2% CA Mann-Whitney U test, (*p* = 0.39, CI: [-64.83, 24.50]). For more information see Extended Data 6-1.

**Table 2.** Composition of the running and stopping task cohorts. A total of 22 mice (11F, 11M) were divided across 2 tasks (running and stopping task) and 2 CA percentages (0% and 2% CA). Starting age corresponds to age on headplate surgery day. Starting weights are from the day before habituation week. No mice were removed from these cohorts.

| Cohort | Sex Composition | Housing Type | Starting Age Range | Average Starting Weights | # of Mice Removed |
| --- | --- | --- | --- | --- | --- |
| Running Task (0% CA) | 3F, 3M | Single-Housed | p60 - p67 | F: 20.6g ± 0.6<br>M: 25.2 g ± 1.3 | 0/6 |
| Running Task (2% CA) | 3F, 3M | Single-Housed | p64 - p67 | F: 21.0g ± 0.7<br>M: 24.4g ± 1.3 | 0/6 |
| Stopping Task (0% CA) | 2F, 3M | Single-Housed | p79 | F: 19.8g ± 0.0<br>M: 27.8g ± 0.7 | 0/5 |
| Stopping Task (2% CA) | 3F, 2M | Single-Housed | p79 | F: 20.2g ± 1.3<br>M: 27.5g ± 0.3 | 0/5 |

We next analyzed peri-event locomotion and licking behaviors from across sessions to investigate event-specific behavioral differences. Consistent with the trend of 0% CA mice traveling further distances, average treadmill speed aligned to RZ entries reveals a speed offset where 0% CA mice generally moved faster than 2% CA mice (Fig. 7*A*). However, a comparison of speeds immediately before and after reward delivery reveals that both groups exhibited similar speed decreases following reward deliveries (Fig. 7*B*). If these speed decreases reflect only reward-associated drinking, similar licking behavior for rewards would be expected. Instead, 2% CA mice exhibited significantly higher lick rates after reward deliveries than 0% CA mice (Fig. 7*C, D,* Extended Data Fig. 7-1). Though, there was no consistent anticipatory licking prior to reward deliveries for either condition. Finally, we examined speed and lick rate aligned to non-rewarded zone (NRZ) entries, different-textured zones that replaced RZs on 10% of trials and were not accompanied by a reward delivery. While there was an initial speed offset between 0% and 2% CA mice as seen with RZs, only 0% CA mice showed a significant speed decrease following zone entry (Fig. 7*E, F*). For NRZ lick rates, we found no consistent pattern of licking in NRZs for either condition (Extended Data Fig. 6-*1C, D*).

**Figure 7.**
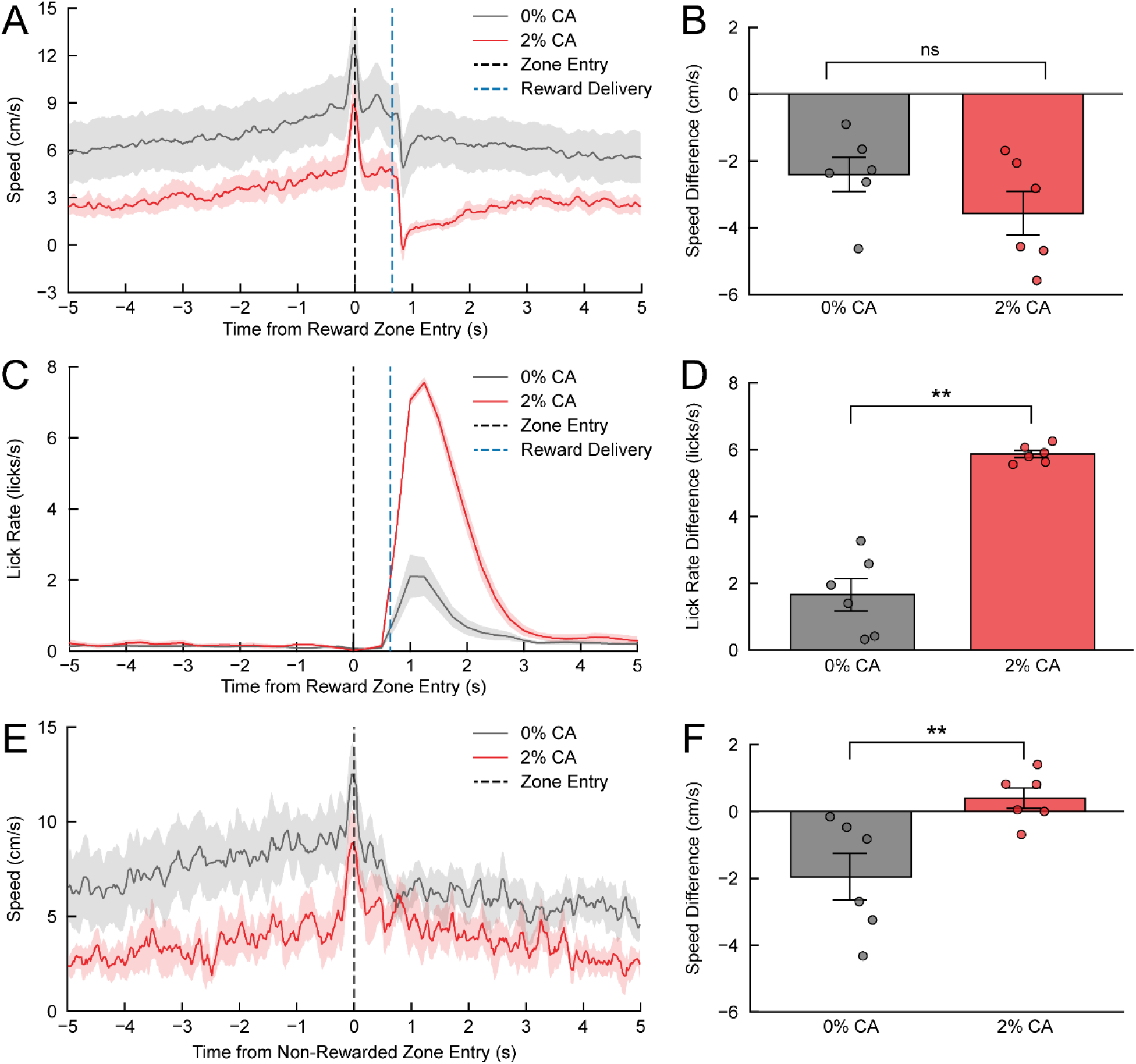
Behavioral epoch analyses for the running task. ***A,*** Event-averaged epoch window for treadmill speed aligned to RZ entries (t = 0) for both 0% and 2% CA mice. Reward deliveries are shown at t = 0.65-s. ***B***, Difference between the 0.65-s to 1.30-s bin and the 0.00-s to 0.65-s bin for both conditions from the RZ-aligned speed plot. Mann-Whitney U test, (*p* = 0.24, CI: [-0.31, 2.95]). ***C,*** Average lick rates in 500-ms bins aligned to RZ entries and smoothed with a 500-ms sliding window. **D,** Difference between the 0.65-s to 1.30-s bin and the 0.00-s to 0.65-s bin for both conditions from the RZ-aligned lick rate plot. Mann-Whitney U test, (*p* = 0.0022, CI: [-5.43, -2.98]). ***E,*** Average treadmill speed aligned to NRZ entry times (t = 0). ***F***, Difference between the 0.00-s to 2.00-s bin and the -2.00-s to 0.00-s bin for both conditions from the NRZ-aligned speed plot. Mann-Whitney U test, (*p* = 0.0087, CI: [-4.21, -0.67]). For more information see Extended Data 6-1 and Extended Data 7-1.

While all these results together are informative regarding the influence of CA water on locomotion-based task motivation, there is a notable limitation of the running task. Rather than indicating decreased task motivation in 2% CA mice, these locomotion and task performance differences could instead reflect that 0% CA mice are motivated for reward-independent running. Therefore, we next assessed a stopping task that rewarded more precise movements rather than distance traveled, as has been similarly demonstrated (Bukwich et al., 2025; Zhang et al., 2021).

### Mice on regular water perform equally as well as mice on 2% CA in a VR stopping task

The second VR task was a stopping task that utilized behavioral shaping to train mice to stop in RZs along a virtual hallway to receive rewards. Because its stopping requirement continually increased in difficulty, this task also assessed how hard mice are willing to work for rewards. Moreover, the stopping task served as motivational control for the running task to help differentiate motivation to run from motivation for rewards. We first evaluated sensitivity (i.e., rewards / reward opportunities) as a metric of performance, because mice could only earn rewards in RZs through correct stopping behaviors. Plotting sensitivity by difficulty level reveals that performance generally decreased across levels for both 0% and 2% CA mice, consistent with the task getting harder (Fig. 8*A*). To quantify sensitivity changes, we analyzed sensitivity across weeks, and while there was no difference in sensitivity by condition, average sensitivity for both conditions monotonically decreased across training weeks (Fig. 8*B*). For another performance metric, we next examined the average rate of level progression for both conditions and found that the rates were linear and almost identical (Fig. 8*C,* Extended Data Fig. 8-1*A*). Given that the difference between difficulty levels is 30 rewards, this finding implies that reward counts remained steady across weeks for both the 0% and 2% CA mice. Consistent with this interpretation, an analysis of reward counts revealed no significant main effect of condition (*F*(1.00, 7.83) = 0.032, *p* = 0.86), training week (*F*(1.23, ∞) = 0.48, *p* = 0.53), or condition-week interaction (*F*(1.23, ∞) = 0.19, *p* = 0.72). Likewise, reward counts analyzed across weekdays showed no condition main effect or condition-week interaction, although there was a significant main effect of weekday (Fig. 8*E*). However, similar to the running task, post hoc comparisons revealed no significant differences between individual weekdays. Together, these findings further indicate that both 0% and 2% CA mice performed the task comparably and maintained consistent reward counts across weeks despite increasing level difficulty. Finally, we also assessed the stopping task for post-reward licking and task acquisition rate as motivational metrics. Consistent with results of the running task, 2% CA mice licked more consistently following reward deliveries (Fig. 8*D*) and initially acquired the task at a rate comparable to 0% CA mice (Fig. 8*F*). Though, one 0% CA mouse never learned to walk on the treadmill and therefore did not progress to level 2.

**Figure 8.**
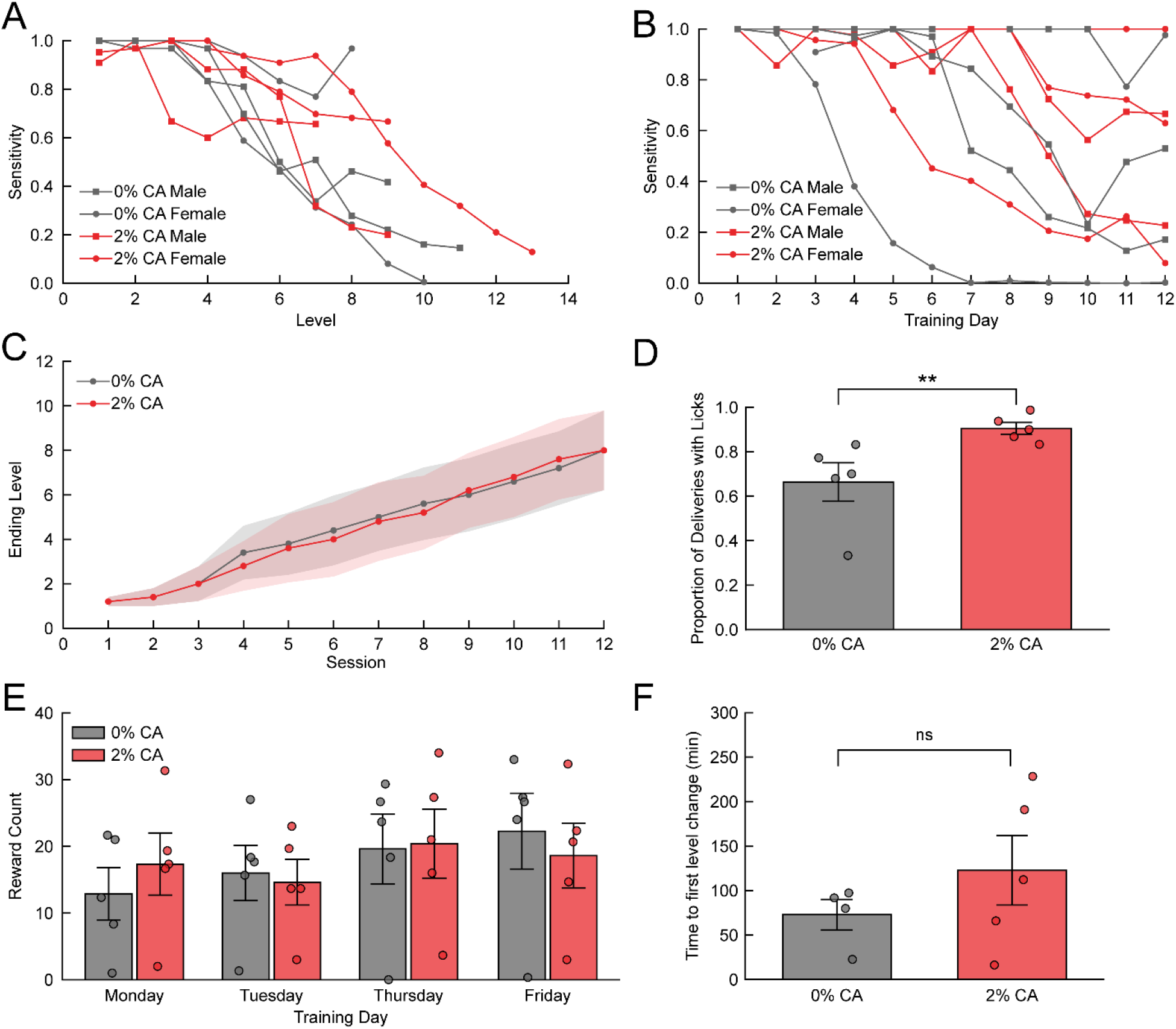
Across-session comparisons for the task performance and engagement results of the stopping task. ***A,*** Level-averaged sensitivity (rewards / reward opportunities) for each mouse from both CA conditions. Individual lines terminate at the final level achieved for each mouse. ***B***, Sensitivity across weeks. Sessions with no RZ trials are excluded from the plot and mice with no RZ trials in any given week are excluded from analyses. F1-LD-F1 test, condition effect (*F*(1.00, 5.85) = 0.89, *p* = 0.38); week effect (*F*(1.67, ∞) = 10.08, *p* = 1.35 × 10^−4^); condition-week interaction (*F*(1.67, ∞) = 0.095, *p* = 0.88). ***C,*** Average rate of level progression for 0% and 2% CA mice. F1-LD-F1 test, condition effect (*F*(1.00, 7.84) = 0.0008, *p* = 0.98); week effect (*F*(1.58, ∞) = 48.04, *p* = 1.42 × 10^−17^); condition-week interaction (*F*(1.58, ∞) = 0.73, *p* = 0.45). ***D,*** Comparison collapsed across sessions of the proportion of reward deliveries that have at least one lick within 2 seconds of delivery. Mann-Whitney U test, (*p* = 0.0079, CI: [-0.53, -0.10]). ***E,*** A breakdown by weekday and CA percentage for the average number of rewards attained on each training day. F1-LD-F1 test, condition effect (*F*(1.00, 8.00) = 0.0054, *p* = 0.94); weekday effect (*F*(1.56, ∞) = 4.83, *p* = 0.014); condition-weekday interaction (*F*(1.56, ∞) = 1.34, *p* = 0.26). Post hoc Wilcoxon signed-rank tests, Mon. vs Tue. (*p* = 1.00, CI: [-4.33, 4.00]); Mon. vs Thu. (*p* = 0.26, CI: [-10.67, 0.83]); Mon. vs Fri. (*p* = 0.16, CI: [-10.67, -0.17]); Tue. vs Thu. (*p* = 0.26, CI: [-10.83, 0.33]); Tue. vs Fri. (*p* = 0.14, CI: [-9.83, -0.33]); and Thu. vs Fri. (*p* = 1.00, CI: [-3.67, 2.33]). ***F,*** Cumulative time across sessions for mice to progress from level 1 to level 2 as a proxy measure for initial task acquisition. Mann-Whitney U test, (*p* = 0.56, CI: [-136.52, 31.36]). For more information see Extended Data 8-1.

Consistent with the running task analyses, we next assessed peri-event locomotion and licking behaviors for CA condition differences in the stopping task. We first visualized average speeds from sample levels aligned to reward delivery events to demonstrate that mice are rewarded only for decreasing speed in RZs (Fig. 9*A*). This also confirmed that levels 9+ required mice to decrease speed to exactly 0 cm/s to earn a reward delivery, as designed. It is important to note that RZs occupy variable distances along the virtual hallway, and mice can slow down anywhere within RZs to get their rewards before they speed up again to leave. Therefore, average speed aligned to RZ entries did not reveal any speed patterns (Extended Data Fig. 8-1*B*). We next quantified post-reward delivery lick rates aligned to reward deliveries and found that 2% CA mice licked significantly more for rewards than 0% CA mice despite earning similar total reward amounts (Fig. 9*B*, *C*, Extended Data Fig. 9-1). Finally, we also compared how 0% and 2% CA mice interacted with NRZs. While 0% CA mice again displayed a trend of slowing down upon NRZ entry, this was not significantly different from the 2% CA mice speed difference (Fig. 9*D, E*). Lastly, neither CA condition exhibited any distinct licking patterns within NRZs (Extended Data 8-1*C, D*).

**Figure 9.**
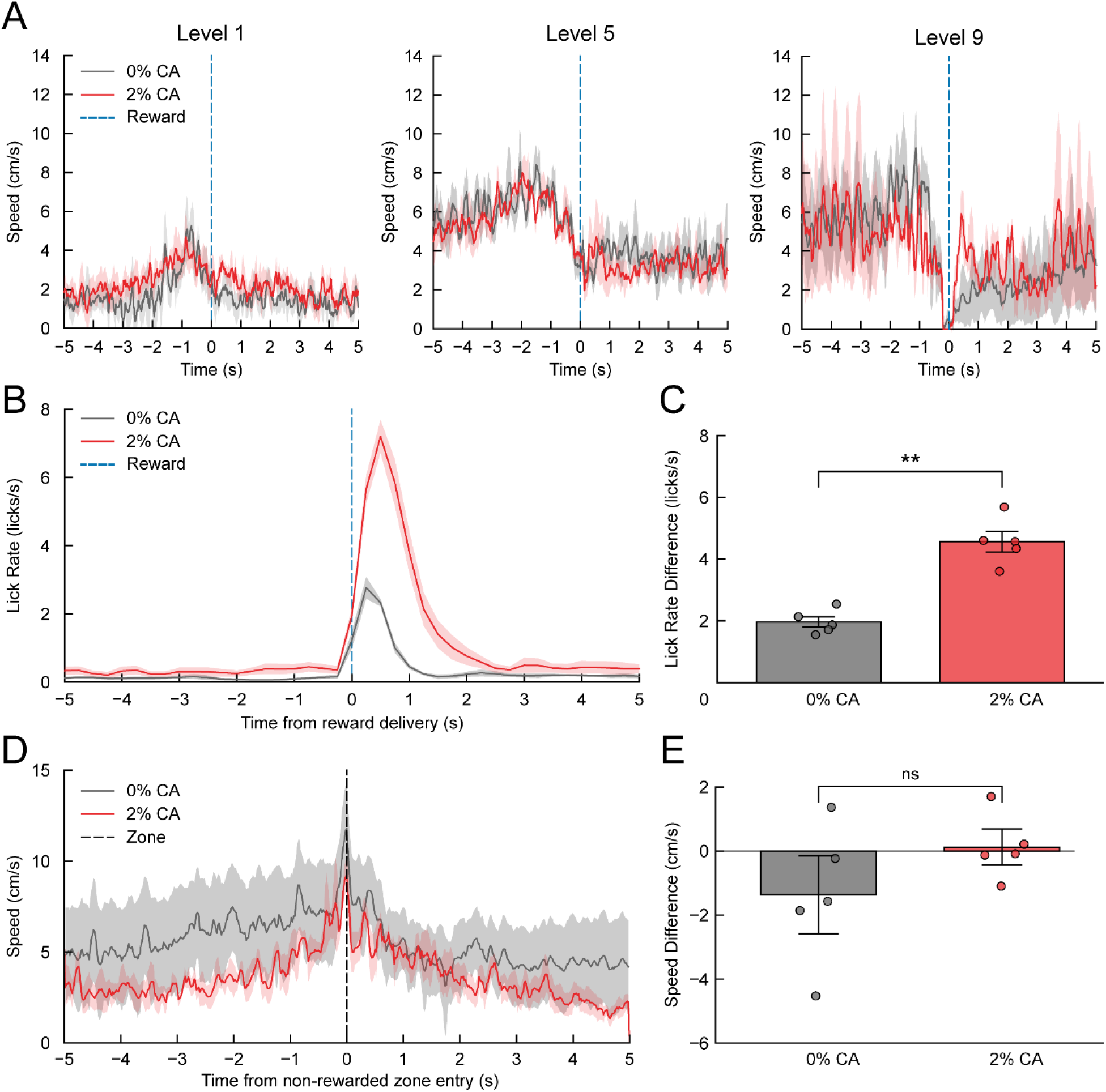
Behavioral epoch analyses for the stopping task. ***A,*** Example speed plots aligned to reward deliveries (t = 0) and averaged for CA condition by level. Level 1 (0% CA, *n* = 5; 2% CA, *n* = 5); Level 5 (0% CA, *n* = 4; 2% CA, *n* = 4); Level 9 (0% CA, *n* = 4; 2% CA, *n* = 3). ***B,*** Event-averaged epoch window for lick rates aligned to reward deliveries (t = 0). ***C***, Difference between the 0.00-s to 0.65-s bin and the -0.65-s to 0.00-s bin for both conditions from the reward delivery-aligned lick rate plot. Mann-Whitney U test, (*p* = 0.0079, CI: [-3.56, -1.89]). ***D,*** Average treadmill speed aligned to NRZ entry times (t = 0). ***E***, Difference between the 0.00-s to 2.00-s bin and the - 2.00-s to 0.00-s bin for both conditions from the NRZ-aligned speed plot. Mann-Whitney U test, (*p* = 0.15, CI: [-3.56, 0.86]). For more information see Extended Data 8-1 and Extended Data 9-1.

## Discussion

We characterized the motivation and health effects of different percentages and schedules of citric acid water by performing daily weight and behavior measurements and by running weekly task-agnostic lick detection sessions. Additionally, we examined the utility of using 2% CA, the most common percentage, as a motivator for two virtual reality treadmill tasks. We examined task performance and mouse motivation across multiple learning and engagement metrics. Our goal is to provide researchers with a more comprehensive overview of the potential suitability of different CA applications such that they can make more informed decisions for their own experiments.

### Slow ramping to 2% CA may maximize motivation and minimize health concerns

In water restriction-type paradigms, total weight change relative to a pre-restriction baseline is the standard metric used to assess animal health (Barkus et al., 2022; Guo et al., 2014; Schwarz et al., 2010; Urai et al., 2021). Using this framework in conjunction with our IACUC guidelines, we found that some CA percentages and schedules resulted in more animal removals than others. However, the number of mice removed from the pair-housed experiments described here overestimates schedule severity due to cage mates being removed along with the mice below the cutoff (even when still above the weight loss threshold). We considered this a necessary limitation as social isolation is a significant stressor (Berry et al., 2012; Rakshasa & Tong, 2020) that could have confounded interpretations on the effects of CA exposure on health and motivation.

Given that all 6 mice in the 4% CA cohort exceeded the 20% weight-loss threshold within three days, we conclude that 4% CA is simply too aversive of an initial concentration. Another schedule that produced unexpected complications was the FR schedule. 1% CA produced comparable initial weight losses between the FR and SR cohorts, but the transition to 2% CA was substantially more challenging for the FR mice. The experiment was terminated after 8 of the 12 mice were removed. We speculate that this outcome resulted from a compounding stressor effect from the environmental instability invoked by rapid increases in citric acid water, consistent with findings from intermittent restriction schedules (Vasilev et al., 2021). Moreover, the extra four days of acclimation provided in the SR schedule may explain why none of the SR mice reached the 20% weight loss cutoff. Therefore, if a ramping schedule is used, additional time between increasing CA concentrations may allow mice to better acclimate to each CA %.

The longitudinal 2% CA and SR cohorts revealed two more successful strategies. The initial weight loss for the 2% CA mice was comparable to the 4% CA mice, but they exhibited a subsequent weight stabilization across time. While 6 of the 12 mice were removed from the study within the first week, the remaining 6 mice all exhibited healthy weight gain as compared to age-matched controls. This suggests that the first week, particularly the first four days, presents the greatest challenge for initial 2% CA exposure. Also consistent with successful adaptation, home cage lethargy generally decreased over time, although week 5 had an increase. Finally, the 2% CA mice exhibited robust lick counts and high frontloading behavior across weeks indicating that long-term habituation to 2% CA water may not decrease the motivation to drink regular water.

In contrast, none of the SR mice were removed despite continued weight loss across the 5 weeks. Although mice on water restriction schedules are generally expected to reach a stable reduced body weight following an initial weight loss period (Schwarz et al., 2010), the SR mice continued to lose weight throughout the experiment. However, this weight loss followed an exponential decay rather than a sustained linear decrease, which indicates that body weight was approaching a stable plateau. These findings suggest that the mice on the SR schedule successfully adapted to the progressive CA concentration increases. Despite this apparent adaptation, the SR mice exhibited more overall anxiety and high lethargy on 4% CA water. Lick detection data indicates there was increased motivational licking from 1% CA to 2% CA but no clear motivational increase for concentrations above 2% CA. We also monitored fecal count, a simple measure that is associated with both stress levels and general hydration (Barkus et al., 2022; Calvo-Torrent et al., 1999; Fertig & Edmonds, 1969). Consistent with habituation, all three of the remaining 5-week cohorts had decreased fecal counts across weeks. However, the SR cohort experienced a significant decrease in fecal count on 3% CA and a potential floor effect not discriminable from typical habituation effects in week 5. While this may be a stochastic pattern, it may also be a physiological indicator of worsening dehydration either from the ramping schedule or the higher CA percentages themselves. Therefore, there is no apparent advantage, and several health disadvantages, to using CA percentages above 2%. Taken together, the most effective balance between animal welfare and experimental utility appears to be a one-step ramp schedule that transitions from 1% to 2% CA after approximately one week. This can be practically achieved by starting mice on 1% CA during habituation procedures and ramping them to 2% CA just prior to task training or as a function of ongoing task performance.

### CA water may not improve running or stopping task performance but may increase reward motivation and decrease behavioral variability

CA water has previously been used to habituate and motivate head-fixed mice to move on treadmills (Jordan et al., 2021; Rupprecht et al., 2024), so we tested the performance differences of mice on 0% and 2% CA during a VR running task. This task crucially relied on mice to initiate their own trials which provided a stronger readout of task motivation (Barkus et al., 2022). Although both groups had similar task acquisition rates and performance improvements over time, 0% CA mice generally traveled farther and progressed to higher levels than 2% CA mice. Superficially, these findings would suggest reduced task motivation in 2% CA mice. However, only the 2% CA mice consistently licked for rewards. Because consummatory licking is frequently considered an essential readout of reward motivation (D’Aquila & Galitsu, 2017), this result implies 2% CA mice are more highly reward motivated despite worse task performance. Therefore, locomotor performance and reward motivation may be partially dissociable under CA water schedules.

One possible explanation for the relatively reduced locomotion in 2% CA mice is that they were conserving water and/or energy. However, studies of water deprivation in rodents report mixed effects on locomotion with reduced water intake producing similar/increased activity (Finger & Reid, 1952; Schwartz et al., 2026; Tucci et al., 2006) or decreased activity (Goltstein et al., 2018; Montgomery, 1953; Rowland, 2007) depending on context. Although the effects of decreased water intake have been studied in wheel running, home cage activity, and maze exploration, the effects on VR locomotion were largely uncharacterized until now. Consequently, it is unclear whether the reduced relative locomotion observed with 2% CA water reflects (1) a reduced physiological capacity to locomote, (2) a reduced motivation to engage in movement, or (3) differences in the relative reinforcing value of running and sucrose, with 0% CA mice potentially exhibiting greater locomotion because running itself has increased hedonic value (Schwartz et al., 2026).

Further evidence of behavioral differences between CA conditions comes from examining non-rewarded zones. Only 0% CA mice exhibited speed reductions within NRZs. One interpretation is that this may reflect texture generalization (Goltstein et al., 2021) and incorrect reward expectation within NRZs, but this is not supported by licking data in NRZs. Though, a lack of licking in NRZs should be interpreted cautiously as neither group exhibited anticipatory licking in RZs. Because this experiment was designed for broader locomotion characterization and not visually guided decision making, we cannot determine if 0% CA mice were worse at texture discrimination. However, a powerful alternative explanation is that 0% CA mice, who enter NRZs at higher speeds on average, slow down as a reflection of distractibility or novelty-seeking (Hughes, 1997; Sarter et al., 2016). This explanation implies that 0% CA mice are less task-engaged than 2% CA mice and may engage in more intrinsically rewarding exploratory strategies.

To determine if the locomotion differences from the running task simply reflect indiscriminate running rather than reward-seeking in 0% CA mice, we next assessed a stopping task. If 0% CA mice are only motivated to run, then they would show limited task progression compared to 2% CA mice because mice only progress in the task by exhibiting correct stopping behaviors. However, 2% and 0% CA mice did not differ in level progression, sensitivity across time, or reward counts. Despite increasing task difficulty and declining sensitivity, reward counts for both CA groups remained stable, which suggests they both adopted successful behavioral strategies and performed more overall trials to offset error rates. As in the running task, however, a key difference was that the 2% CA mice exhibited more consistent reward-licking, which we interpret as higher reward motivation. Taken together, these findings indicate that task performance alone does not fully capture behavioral differences between CA groups and instead imply subtler differences in motivation or behavioral strategy.

Notably, one 0% CA mouse stopped performing the stopping task and started continuously running after the full-stop (0 cm/s) requirement was introduced. Although based on a single mouse, this behavioral shift to a more immediately rewarding behavior is consistent with the notion that locomotion itself can be rewarding. This would imply that 0% CA mice are more likely to abandon the task when the task is difficult, which is corroborated by how more difficult tasks require more than just sweetened rewards to motivate non-water restricted mice (Bramati et al., 2023). However, because the remaining 0% CA mice did not adopt this strategy, hedonic running alone cannot explain all CA group differences. Likewise, NRZ data reveals that only 0% CA mice had a trend of slowing down in these zones, but this was not significantly different than the 2% CA mice. This trend is consistent with the interpretation that 0% CA mice exhibit greater exploratory or information-seeking behaviors. The decrease may have been less drastic than seen in the running task because the stopping task required more varied movement speeds. Thus, while 0% CA mice are still reward motivated, they may be more distractible and/or susceptible to other intrinsically motivating behaviors that are not directly related to task goals. Therefore, the results of both VR tasks together suggest CA water may reduce the natural variability of mouse locomotor behaviors by controlling for other competing motivations. CA did not improve task performance but did produce behavioral readouts that were more readily attributable to reward motivation.

### Limitations and further considerations

While we have expanded the understanding regarding applicability of citric acid water, we encourage future investigations into the uses of half-percentages and performance-based ramping schedules, which are other techniques currently in use (Mai et al., 2024). For our characterization cohorts, we measured motivation in a task-agnostic way via water licking to broaden the relevance of our measure, which comes at the cost of not directly knowing how motivation will transfer to any specific task. Additionally, the results of our 2% CA characterization cohort should be interpreted cautiously as we could only study mice who successfully adjusted and did not cross the weight loss cutoff.

The findings from our VR cohorts also have limitations to consider. Sample sizes were intentionally kept small to minimize the number of mice undergoing headplate surgeries, but this reduced our power to detect more subtle differences between CA groups. Additionally, our findings may not readily transfer to freely roaming or non-VR movement-based tasks due to the potential of the VR system changing how mice engage with their surroundings. Our tasks were designed to give mice individual freedom in how they engaged with the VR system for the purpose of broad behavioral characterization. Given that we did not find the stark differences between 0% and 2% CA mice that we expected, longer training schedules or punishments for non-task related behaviors may help further disambiguate the behaviors and motivations of the two groups. A further consideration is that we selected 5% sucrose as our reward to serve as a middle ground between complex behavioral paradigms that use only water rewards (Kondo et al., 2025; Mai et al., 2024) and those that use highly sweetened rewards (i.e., 10% - 30% sugar) (Rupprecht et al., 2024; Urai et al., 2021). Regular water would likely have produced starker differences between 0% and 2% CA mice while more highly sweetened water might have reduced the differences.

Despite having limitations, our findings do raise questions about whether certain VR movement-based tasks need additional external motivational factors beyond a sweetened reward. Furthermore, if the goal is for mice to run or patch-forage, which is the ethologically relevant behavior underlying both of our tasks (Oesch et al., 2024), then a water restriction or CA schedule may not be necessary. Other researchers have been considering alternative motivation methods for other task types that do not involve water restriction at all (Grayson et al., 2025; Ma et al., 2023). Moreover, CA water may limit the natural variability in mouse behavior by restricting competing motivations. Thus, experimenters should carefully weigh the benefit of increased behavioral consistency against the cost of decreased information regarding naturally occurring behavioral variation. A major rationale for using CA water over traditional water restriction is to improve animal welfare while maintaining experimental rigor (Urai et al., 2021). This consideration encompasses both animal wellbeing and the generalizability of behavioral results. Therefore, we believe more research needs to be conducted into understanding and improving motivation for complex tasks, especially VR-based locomotion tasks. This research would further clarify the extent to which water restriction-type paradigms are necessary for achieving reliable performance in these tasks, and whether alternative motivational strategies can produce equivalent behavioral performance while further improving animal welfare.

## Supporting information

Statistics Table

---

**Extended Data 2-1.**
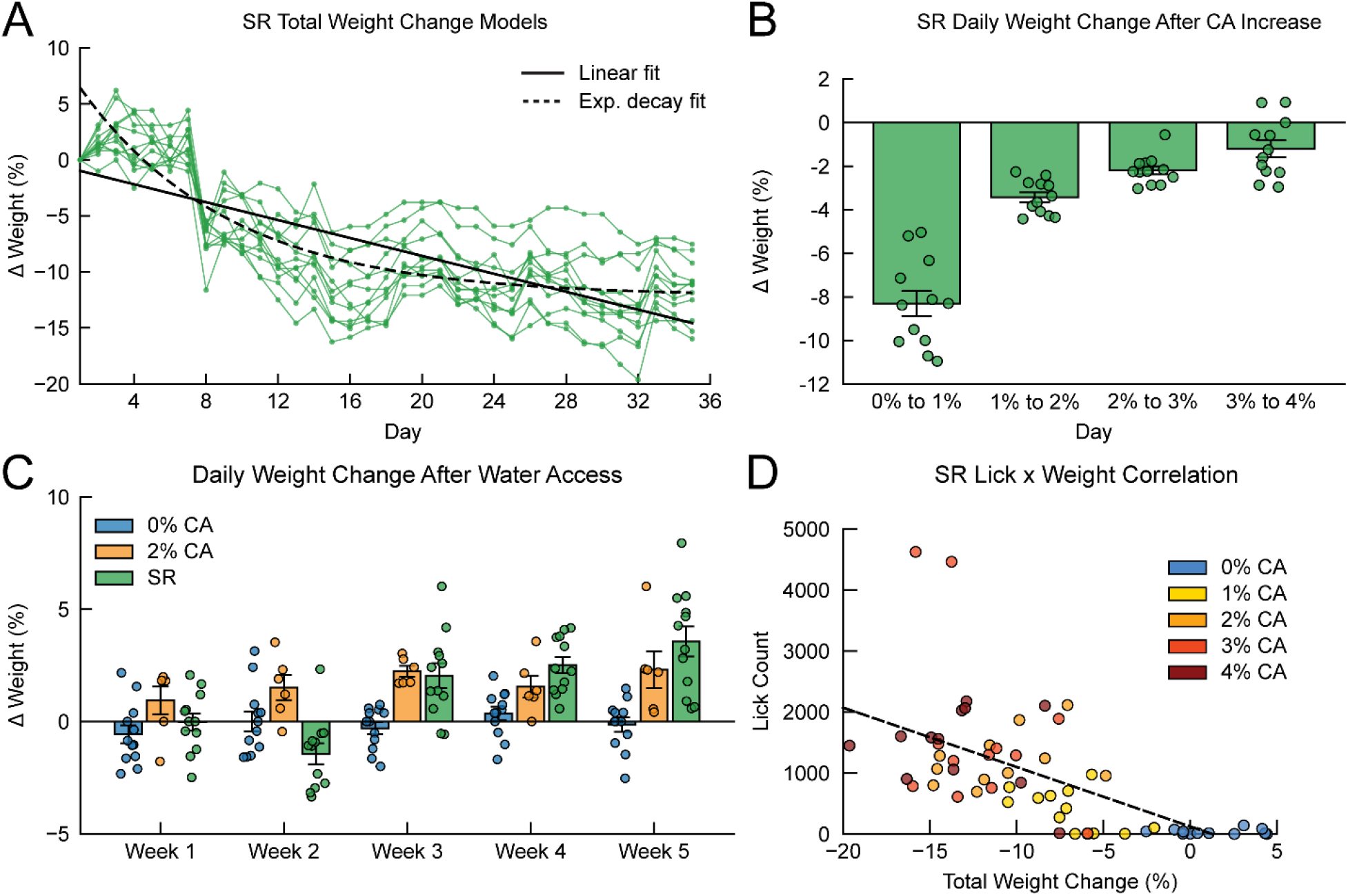
Further examination of weight metrics for the characterization cohorts. ***A***, Model comparison for the SR total weight change across time normalized to baseline weights. Linear model (Marginal R^2^ excluding random intercept variance: 0.53, AIC: 2166.65) vs exponential decay model (Pseudo-R^2^: 0.79, AIC: 1960.94). ***B,*** SR relative weight changes following successive increases of CA concentration. Friedman test, (χ²(3) = 31.42, *p* = 6.92 × 10^−7^). Post hoc Conover-Iman tests, 0% to 1% CA vs 1% to 2% CA (*t*(33) = -5.57, *p* = 6.93 × 10^−6^, CI: [-6.41, -3.28]); 1% to 2% CA vs 2% to 3% CA (*t*(33) = -6.42, *p* = 8.39 × 10^−7^, CI: [-2.14, -0.59]); and 2% to 3% CA vs 3% to 4% CA (*t*(33) = -1.28, *p* = 0.21, CI: [-2.28, 0.10]). ***C,*** Relative weight changes by 0% CA, 2% CA, and SR cohorts on the days after 30 minutes of access to water. F1-LD-F1 test, cohort effect (*F*(1.98, 20.51) = 38.56, *p* = 1.19 × 10^−7^); week effect (*F*(3.13, ∞) = 7.80, *p* = 2.42 × 10^−5^); and cohort-week interaction (*F*(4.98, ∞) = 4.41, *p* = 5.26×10^−4^). ***D,*** Same repeated measures correlation as Fig. 4H but color coded by CA percentage.

**Extended Data 5-1.**
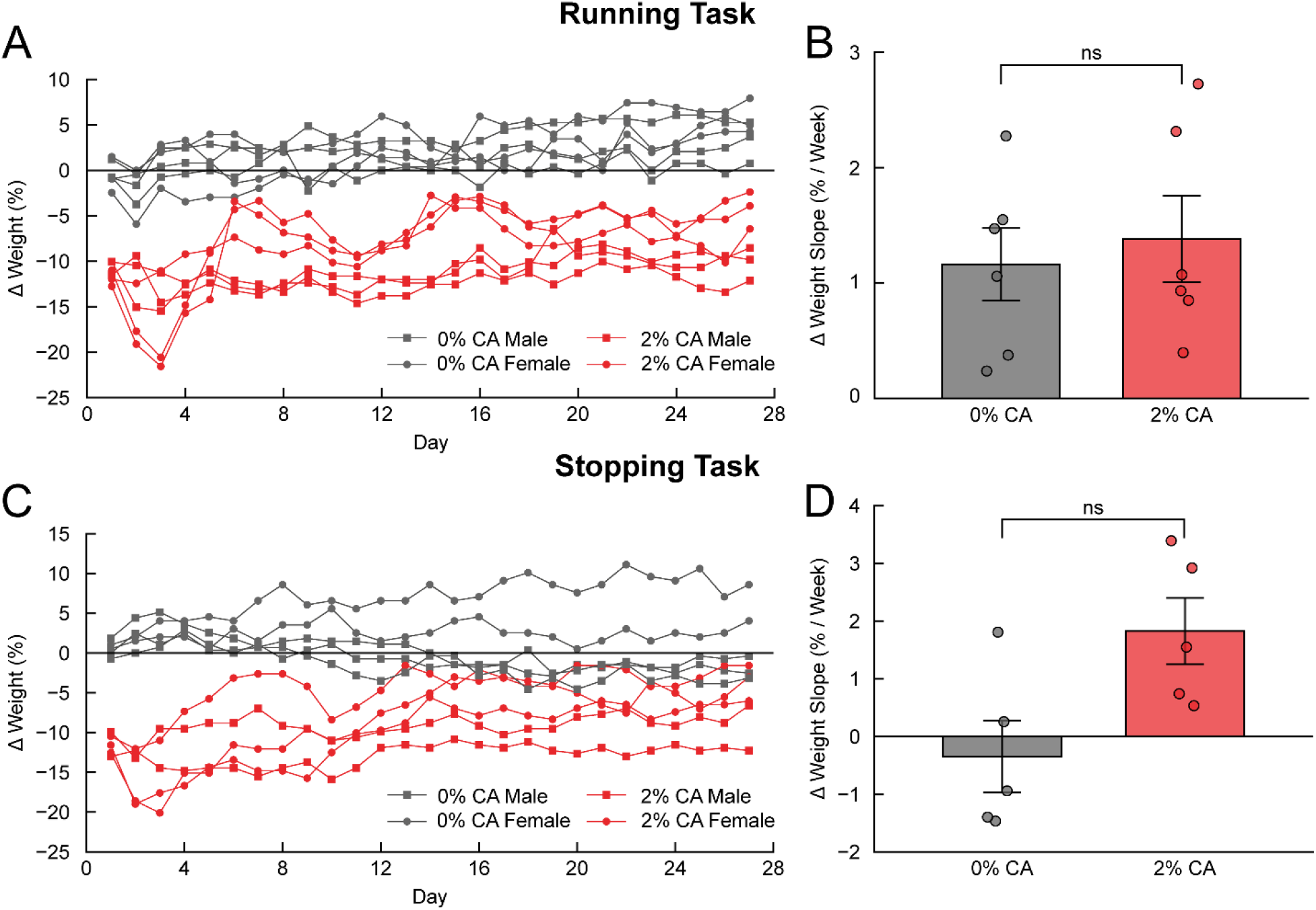
Running and stopping task cumulative weight changes across habituation week and training weeks. ***A,*** Weight fluctuations compared to baseline weights of the running task mice. ***B***, Rate of weight change comparison for both CA conditions of the running task. Mann-Whitney U test, (*p* = 0.70, CI: [-1.18, 0.70]). ***C,*** Weight fluctuations compared to baseline weights of the stopping task mice. ***D***, Rate of weight change comparison for CA both conditions of the stopping task. Mann-Whitney U test, (*p* = 0.056, CI: [-4.32, -0.48]).

**Extended Data 6-1.**
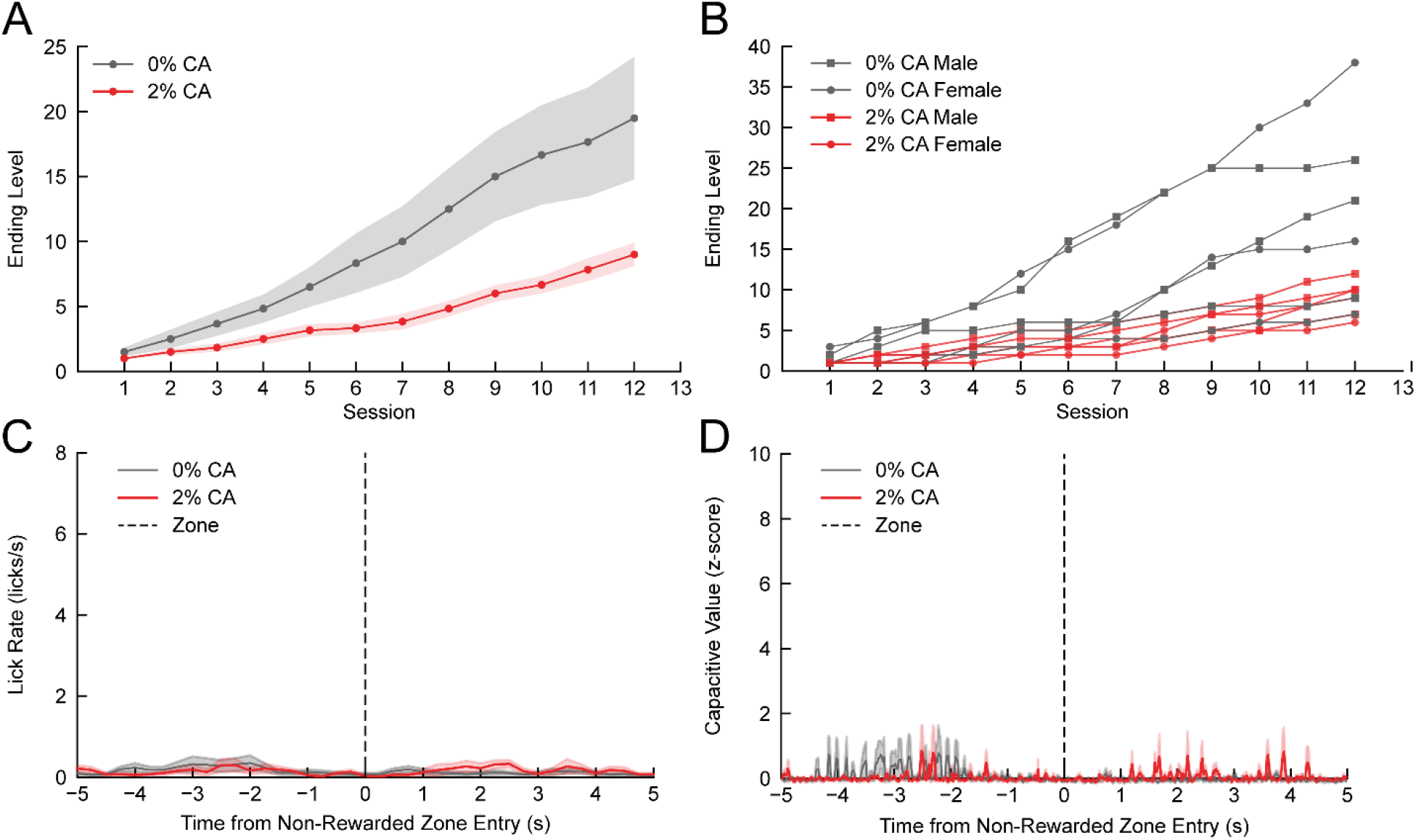
Further comparisons of the 0% and 2% CA mice of the running task. ***A,*** Average rate of level progression by CA condition. F1-LD-F1 test, condition effect (*F*(1.00, 8.61) = 7.05, *p* = 0.027); week effect (*F*(1.41, ∞) = 147.04, *p* = 2.36 × 10^−46^); and condition-week interaction (*F*(1.41, ∞) = 3.95, *p* = 0.033). ***B,*** Rate of level progression for individual mice. ***C,*** Average lick rates aligned to NRZ entry (t = 0). ***D***, Average capacitance traces aligned to NRZ entry (t = 0).

**Extended Data 7-1.**
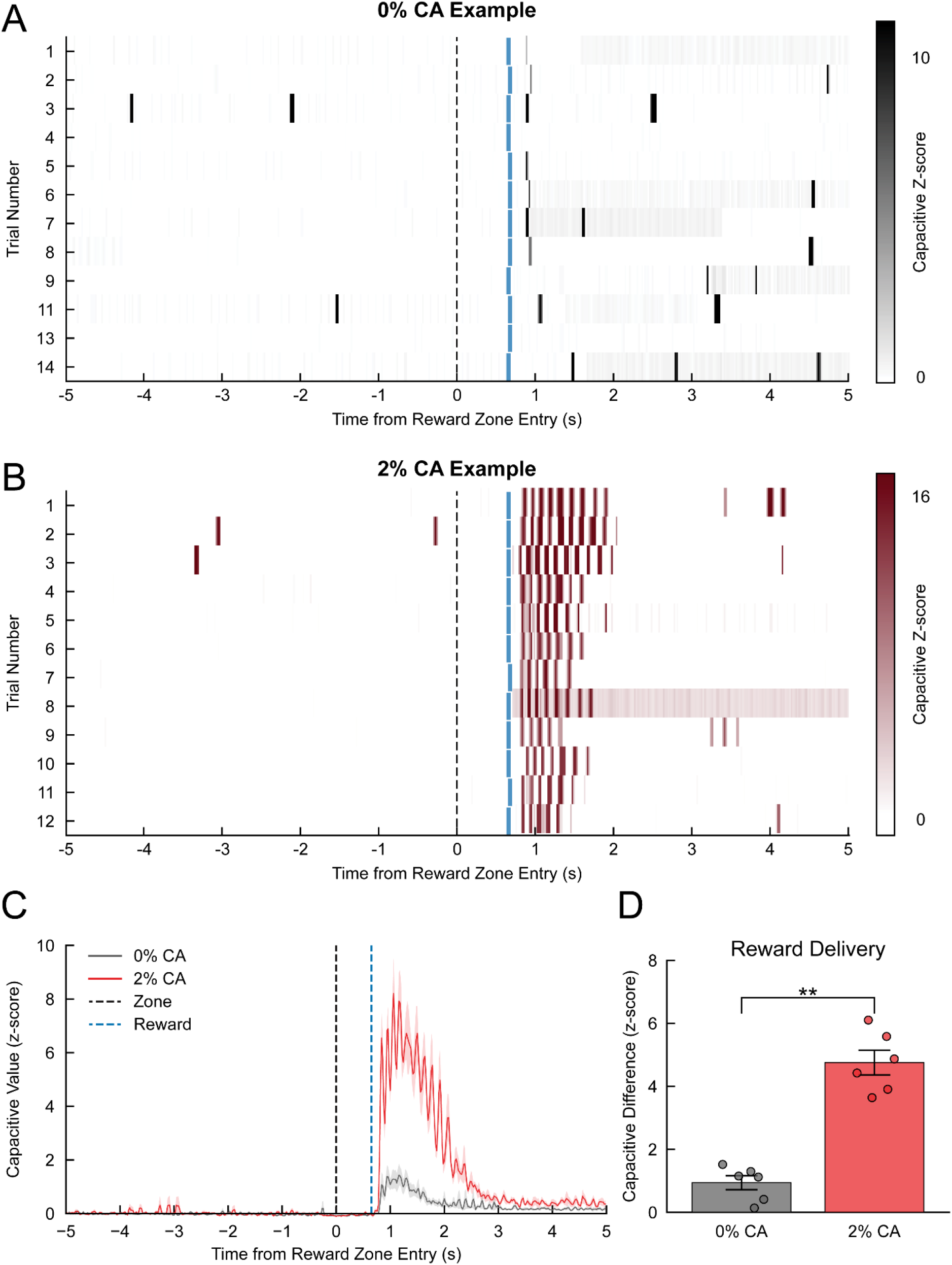
Further licking analyses for the running task. ***A,*** Sample capacitance lick raster plot from session 12 for a 0% CA female mouse. ***B,*** Sample capacitance lick raster plot from session 12 for a 2% CA female mouse. Both ***A*** and ***B*** depict trial numbers as including any NRZ trials, which are not shown. All trials are aligned to the point of RZ entry (t = 0), and blue vertical bars indicate reward deliveries (t = 0.65). ***C,*** Event-averaged epoch window for capacitance aligned to reward deliveries (t = 0) as a proxy for licking behavior. ***D***, Difference between the 0.65-s to 1.30-s bin and the 0.00-s to 0.65-s bin for both conditions from the RZ-aligned capacitance plot. Mann-Whitney U test, (*p* = 0.0022, CI: [-4.77, -2.76]).

**Extended Data 8-1.**
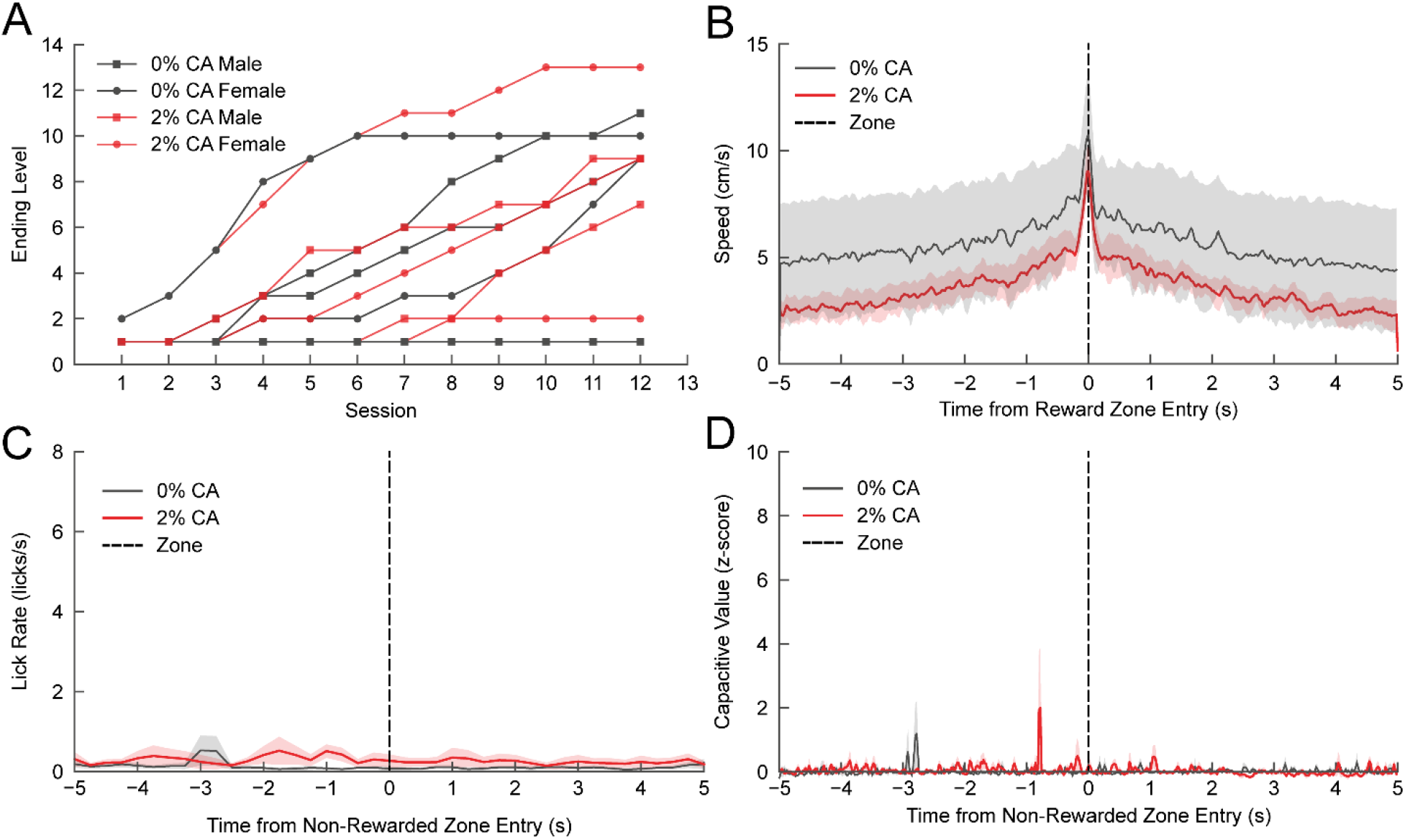
Further comparisons of the 0% and 2% CA mice of the stopping task. ***A,*** Same as Fig. 8C but showing individual traces per animal. ***B,*** Average speeds aligned to reward zone entries (t = 0). ***C,*** Average lick rates aligned to NRZ entry (t = 0). ***D***, Average capacitance traces aligned to NRZ entry (t = 0).

**Extended Data 9-1.**
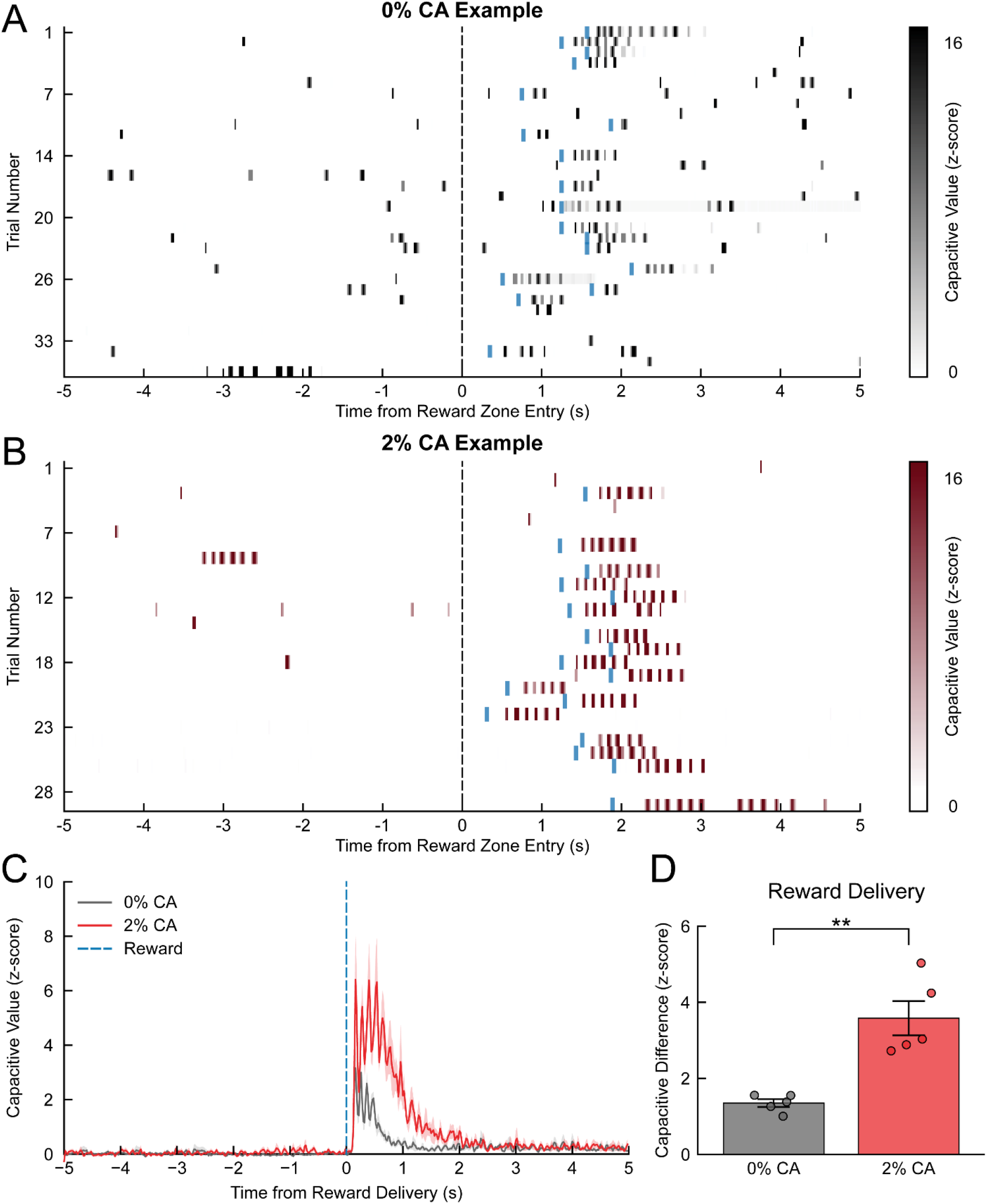
Further licking analyses for the stopping task. ***A,*** Sample capacitance lick raster plot from session 12 for a 0% CA male mouse. ***B,*** Sample capacitance lick raster plot from session 12 for a 2% CA female mouse. Both ***A*** and ***B*** depict trials numbers as including any NRZ trials, which are not shown. All trials are aligned to the point of RZ entry (t = 0), and blue vertical bars indicate reward deliveries at variable times depending on reward delays and stopping behaviors. ***C,*** Event-averaged epoch window for capacitance aligned to reward deliveries (t = 0) as a proxy for licking behavior. ***D***, Difference between the 0.00-s to 0.65-s bin and the -0.65-s to 0.00-s bin for both conditions from the reward delivery-aligned capacitance plot. Mann-Whitney U test, (*p* = 0.0079, CI: [-3.48, -1.34])

## Methods

### Subjects

All experiments performed within this study were conducted in accordance with [Author University]’s Institutional Animal Care and Use Committee (IACUC). Female (*n* = 40) and male (*n* = 36) C57BL/6J mice (#000664, Jackson Labs) ages P60-123 were generated through in-house breeding (see Table 1 and Table 2). All mice were provided with *ad-libitum* food access (LabDiet® 5053 - PicoLab® Rodent Diet 20) for the duration of the study and were housed in reverse light/dark rooms (ZT12- lights off at 10:00 AM) held at ∼69°F and ∼45% humidity for at least a week prior to and throughout experimental testing. All weight and home cage measurements occurred within ZT12 and ZT14 while behavioral tasks were performed between ZT14 and ZT18.

### Citric acid water

The citric acid (Sigma-Aldrich, Product # C0759) water prepared for all experiments used a mass/volume formula (i.e., 2% = 2g of CA in 100 mL home cage water) as previously reported (Urai et al., 2021). CA was thoroughly mixed into water until fully dissolved. Bottle changes occurred weekly on Saturdays (unless otherwise specified) and were performed only after daily measurements. To maintain measurement consistency, the same experimenter handled all mice, recorded all behavioral data, and mixed all CA water for all cohorts.

### Characterization cohorts

#### CA SCHEDULING

For the 4% CA cohort (Table 1), triple-housed mice (3F, 3M) were given home cage water bottles containing 4% CA water and weighed daily. This cohort was a smaller pilot cohort run prior to the pair-housed 5-week characterization cohorts. Given the results of this pilot, a 5-week characterization for a larger pair-housed 4% CA cohort was not examined. For the longer 5-week characterization experiments, mice (26F, 22M) were sorted into 4 separate cohorts: 0% CA, 2% CA, fast ramp (FR), and slow ramp (SR) (Table 1). Mice were first pair-housed by removing additional siblings, and then an equal number of female and male cages were randomly assigned to each cohort. For the duration of the characterization experiments, each cohort followed a set CA schedule (Fig. 1). The 0% CA cohort was always on regular water, and the 2% CA cohort was always on 2% CA water. The SR cohort started on regular water and successively moved to a higher CA percentage every week. This resulted in them having a full week of use for each percentage between 0% and 4% CA. The FR cohort also followed a ramping schedule, but moved to a higher CA percentage every 3 days instead of every week (i.e., bottles were changed every 3 days). When the FR mice reached 4% CA water on Day 13, the experiment was designed for them to stay on 4% CA for another 3 weeks.

#### WEIGHT AND BEHAVIORAL DATA

All mice were weighed daily and removed from the study if they lost more than 20% of their starting weight per our IACUC guidelines. Mice were also assessed daily on 3 qualitative home cage behavioral metrics: nest making, anxiety, and lethargy. Nestlets were replaced weekly during cage changes. During daily checks, cages of mice were scored as having no nest if the paper nestlet was less than 50% shredded/dissociated. Individual mice were scored as being lethargic if they barely moved inside the home cage, made no attempt to escape experimenter grasp, or did not exhibit rearing and/or circling behaviors inside the weigh cup. Individual mice were scored as being anxious if they were seen frantically digging through their bedding or running laps around the home cage/trying to escape the cage. For each behavioral metric, the presence of any qualifying observation on a given day resulted in that day being counted as an aberrant behavior day of that behavioral metric. Weekly aberrant behavior percentages were then quantified for each metric by calculating the percentage of observation days on which each behavior was present (i.e., 3 anxious days/ 7 total days = ∼43% anxious observations for that week).

#### 24-CAGE LICKOMETER SETUP

Lick detection was performed using a custom 24-cage lickometer setup comprised of two daisy-chained capacitive touch sensor boards (MPR121, Adafruit^®^) connected to a microcontroller (Arduino^®^ UNO) running a custom script (Fig. 4A). Prior to the start of experiments, mice were separated into empty cages for two 30-minute habituation sessions to the cages and soldered bottle spouts (Ext. Data Video 1). The cages were empty to allow for easier counting of fecal pellets and for the ease of cleaning and reusing the cages across weeks. Capacitance changes of the bottle spouts, as detected by the microcontroller script, were recorded using a Raspberry Pi^®^ 5 running a custom Python logging script (see [Author’s GitHub page]). All mice licked for water at least once across both habituation sessions which indicates that any later sessions with 0 licks are not a result of the mice having neophobia of the spouts. During the weekly experimental sessions, mice were again separated into empty cages and given 30 minutes of access to regular water. This occurred every Wednesday for the 0% CA, 2% CA, and SR cohorts while the FR cohort was scheduled to have their water access sessions on only their last 3 Wednesdays. At the end of each session, mice were returned to their home cages and their fecal pellets left in the empty cages counted and recorded.

### Virtual reality (VR) cohorts

#### HEADPLATE SURGERIES

For all VR mice (11F, 11M), headplate surgeries were performed using isoflurane in oxygen as the anesthetic and extended-release buprenorphine (3.25 mg/kg, Ethiqa XR^®^) as the pre-operative analgesic. Mice were first induced at 3% isoflurane and maintained at 1-2% for the duration of the surgery. Briefly, the scalp was removed and the skull scored with a dental drill (BP50, Ram Products Inc.) to improve dental cement adhesion. Custom titanium headplates were attached and exposed skull covered using dental cement (Metabond^®^, Parkell). After surgeries, mice were placed in single-housed cages to mitigate damage to healing skin by cage mates. Supplemental post-operative pain management was provided as needed via subcutaneous injections of meloxicam (5.0 mg/kg). Wooden chew bars were provided for additional enrichment, and mice were allowed to recover for a week with access to DietGel^®^ boost (ClearH_2_O^®^) as a dietary supplement.

#### HABITUATION PROCEDURES

Following one week of surgery recovery, mice underwent one week of habituation procedures. Within each of the two VR cohorts, half of the mice were randomly assigned to receive 2% CA water while the other half continued on regular water (0% CA) (Fig. 5*B*). Any 2% CA mice that initially exceeded 20% weight loss were provided one hour of access to regular water before being returned to 2% CA water.

Experimental habituation consisted of five consecutive days during which mice had the opportunity to consume up to 1 mL of 5% sucrose water each day. The first two days consisted of handling habituation in which mice were gently cupped and syringe-fed sucrose water until they became comfortable with being held and handled. The next two days were 15 and 30 minutes of treadmill head-fixation habituation. During these sessions, mice were encouraged to walk forward on the treadmills by using the sucrose water as a scent lure and by slowly rolling the tread backward to encourage walking. For the last day of habituation, mice were head-fixed for 30 minutes with the treadmills now placed inside of the behavior boxes. As such, their sucrose water was delivered from the reward spouts in the behavior boxes (Fig. 5*A*).

#### BEHAVIOR BOXES AND TREADMILLS

The behavior boxes were custom built using Thorlabs^®^ equipment for the metal base plates and structural features. The boxes had two monitors (Miktver^®^, 10.5” IPS LCD, 1920×1280 FHD, 16:10 aspect ratio) placed ∼4.5” from the eyes of the mice (Fig. 5*A*). The monitors were placed together at a ∼120 angle to simulate the corridors of a hallway on either side of the mice. Just below the monitors was a reward delivery spout (20G blunt tip needle) soldered to be part of a capacitive circuit. This setup used the Arduino^®^ CapacitiveSensor library with one Arduino^®^ Mega and circuit per behavior box. Sucrose water (5%) for each box was contained within 60 mL reservoirs and connected through tubing to the delivery spouts. The system was gravity fed and controlled by a solenoid valve (MB2029, Gems Sensors & Controls^®^) that could be triggered manually with a button (during habituation and spout positioning prior to each training session) or by control from the virtual hallway program running on each behavior box’s computer. Delivery amounts were back-calculated using a custom solenoid calibration script that converted solenoid opening time into desired volume delivery by accounting for gravity-fed pressure. Calibrations were performed separately for each box just prior to the start of the VR experiments. The linear treadmill models used were modified from the Janelia rodent-belt-treadmill model (Arnold, 2023) and had smaller base plates that allowed them to easily be moved in and out of the boxes. Each treadmill had an encoder (HEDR-5420-ES214, Broadcom Limited^®^) that was monitored by an Arduino^®^ Teensy board running the Janelia-provided treadmill script further modified for use by our lab (see [Author’s GitHub]). This script allowed for the recording of accurate distances and speeds achieved by the mice during their VR training sessions.

#### VR RUNNING AND STOPPING TASKS

The closed-loop, endlessly rendering virtual hallway used for the running and stopping tasks was developed by our lab and can be found in detail on [Author’s GitHub]. Briefly, the VR hallway was built using the Panda3D game engine (Goslin & Mine, 2004) and implements a level-based training system in which an experimenter can define numerous task parameters for each level. While the two tasks differed on these training parameters, the basic functions of the hallway were the same. Initially, the hallway displayed a default wall texture for a pseudorandom distance sampled from a discrete distribution determined by the current level. Once a mouse walked the predetermined distance on the treadmill, all walls changed to either the reward zone texture (RZ; 90% probability) or the non-rewarded zone texture (NRZ; 10% probability), thus marking trial onset (Ext. Data Video 2, Ext. Data Video 3). In both tasks, mice could earn a 3µL delivery of 5% sucrose water in RZs, although reward delivery depended on task-specific contingencies. To return the walls to the default hallway texture and initiate the next inter-trial interval, mice had to travel a second, shorter pseudorandom distance. Thus, both the running and stopping tasks required voluntary locomotion to initiate trials and determine the pace of training, while the use of pseudorandom lengths reduced temporal predictability. In real time, a TCP socket server monitored the performance of the mouse (i.e., reward counts) and progressed the task to more difficult levels (i.e., more challenging task parameters) within the same session and while maintaining progress across multiple sessions. This enabled mice to engage in self-paced training in which task parameters evolved according to each individual mouse’s performance.

For the running task, every RZ entry was followed by a reward after a 650ms delay, regardless of whether the mouse remained within the RZ (Fig. 5*C*). Mice automatically progressed to the next difficulty level every 15 rewards. At each successive level, the average inter-trial distance (i.e., the length of the default-textured hallway between trials) increased exponentially, thus requiring mice to travel farther distances to keep earning rewards. Based on previous data collected in VR hallways, the inter-trial distance was increased gradually through small increments rather than large steps. In the VR system, all distances are defined by the number of wall segments traversed with one wall segment equivalent to an actual distance of ∼3.0 cm. Accordingly, the mean number of wall segments used to generate the pseudorandom distribution of inter-trial distances increased according to a slowly growing exponential function. The mean number of inter-trial wall segments (*y*) for each difficulty level (*x*) was calculated as:

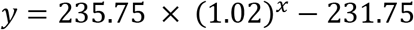

Like the running task, the stopping task initially rewarded mice just for walking to RZs (Fig. 5*C*). However, the inter-trial distances did not continually increase and the threshold for level progression was 30 rewards (versus 15 in the running task). Instead, mice were trained to stop walking within RZs using three different training parameters: reward delay, reward zone size, and zero-speed detection. The first 8 difficulty levels increased the reward delay (i.e., threshold of time spent within the RZ to earn a reward). Therefore, mice were required to spend increasingly more time inside of RZs to get rewards and would not receive the reward if they left before the reward delay was reached. Additionally, the average size of RZs progressively decreased as difficulty levels increased such that mice needed to stop moving more quickly upon entering a RZ to remain within the RZ. Levels 9 + of the training were the most difficult as they required mice to fully stop moving (0 cm/s) for increasingly longer periods of time within RZs to receive rewards.

### Data processing and analyses

All data visualizations and analyses were conducted using custom Python scripts that implemented both Python and R statistical packages and are documented in detail on [Author’s GitHub]. In brief, Python packages for analyses included NumPy, pandas, matplotlib, SciPy, pingouin, statsmodels, and rpy2 (Gautier, 2026; Harris et al., 2020; Hunter, 2007; Seabold & Perktold, 2015; The pandas development team, 2026; Vallat, 2018; Virtanen et al., 2020).

#### 24-CAGE LICKOMETER DATA

The lickometer capacitive data from the 5-week characterization cohorts were first visually inspected for quality control. Next, capacitive data for each sensor were normalized to a baseline calculated as the peak of the Gaussian kernel density estimation (KDE). Then, the first-order derivative was calculated using a forward difference to these normalized values and peaks were extracted using a peak detection algorithm set for values above the arbitrary threshold 0.1. These peaks were defined as licks, and quantifications were robust despite capacitive interference caused by mice grasping the spout with forelimbs.

#### VR DATA

For VR-specific pre-processing, all behavioral data were also first scanned for quality control. Capacitive recordings with major baseline artifacts (4 files) and treadmill files with no recorded data due to encoder problems (3 files) were removed. The remaining files were batch cleaned to remove Arduino^®^ startup artifacts and minor sensor baseline artifacts. The lick detection algorithm for the capacitance data first utilized a KDE peak normalization. Under the assumption that capacitive readings followed a bimodal distribution, a dynamic threshold was calculated by fitting a second Gaussian KDE to the normalized data. This allowed the use of a ‘deepest valley’ algorithm to find the gap between the peak of the baseline and the peak of true lick signals. A minimum value guard and a post-valley signal guard were employed to ensure sessions with few licks would be processed accurately. We then applied a peak detection function to values above the computed threshold, and the peaks were defined as licks.

For VR-specific epoch analyses, peri-event time series of treadmill, lick rate, and capacitance data were aligned to zone entries or reward deliveries. Event histories were first reconstructed by matching trial log entries across data streams for each session and animal. Raw treadmill speed and capacitance traces were resampled at their native 50 Hz sampling rate, and capacitance signals were z-scored within each session. Trials containing capacitance recording gaps (i.e., removed artifacts) within the epoch window were excluded from further analyses. For each event, a window was extracted from the uniformly sampled signals and interpolated onto a common time axis. Lick rate was calculated in 500-ms bins using the VR lick detection algorithm and smoothed with a 500-ms sliding window. Epoch averages were computed hierarchically with trial traces averaged within sessions, session means averaged within mice, and mouse means averaged within experimental conditions. For level-stratified analyses, data were first concatenated before being sliced into levels based on the timestamp for the final reward in each level. The difference bar plots were generated by partitioning each peri-event epoch into predefined sub-windows. The sub-window sizes were determined based on the event of interest, and the pre-window and post-window were kept the same size. For the running task RZ speed and lick rate analyses, 0.65-s bins were used to encompass the interval between zone entry and reward delivery while the NRZ zone speed analysis used 2-s bins to adequately capture longer timescale locomotion changes. For consistency, the same window sizes were used for all stopping task analyses. Signal averages were calculated within each sub-window before subtracting the pre-event window values from the post-event window values.

#### WEIGHT DATA

Weight changes for all cohorts were calculated as a total percentage change from baseline and a daily percentage change between consecutive measurements using the following equations:

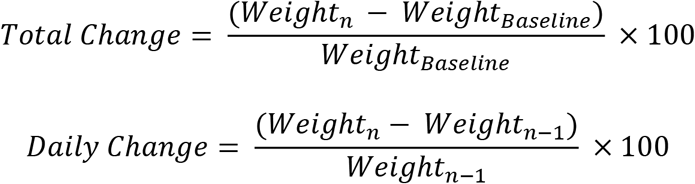

Baseline weights for the total change calculations were obtained from a pre-experiment weighing session (day 0) except for the two ramping cohorts, for which the baseline was calculated as the mean weight across all 0% CA days. However, to facilitate comparisons between the SR and the 0% and 2% CA cohorts, total weight change for the SR cohort was recalculated using day 1 as the baseline. Consequently, day 1 total weight change values were fixed at 0% change, so this day was excluded from weekly averages. Total changes for the FR cohort were not re-calculated because the experiment was terminated before completion due to animals having crossed the IACUC approved weight-loss thresholds. Weekly rates of total weight change were found by fitting individual linear regression lines to each animal’s weekly average total weight change. Additional SR cohort slope fitting was conducted using mixed effects linear and exponential decay fits with random intercepts for individuals using R packages lmerTest, nlme, and lme4 (Bates et al., 2015; Kuznetsova et al., 2017; Pinheiro et al., 2026).

#### STATISTICAL ANALYSES

For statistical tests, we assessed all data for the Gauss-Markov assumptions of ordinary least squares (OLS) estimation models using Python packages and R packages MASS, car, and fitdistrplus (Delignette-Muller & Dutang, 2015; Fox & Weisberg, 2019; Venables & Ripley, 2002). Given the number of outliers and OLS assumptions violations, we used nonparametric tests for much of the dataset. We report main findings within figure legends and the results section, but additional analyses and further details (i.e., sample sizes, effects sizes, etc.) can be found in the stats table (Table 3). Nonparametric repeated-measures tests were conducted using base R v4.6.0 and the nonparLD package (Noguchi et al., 2012; R Core Team, 2026). Factorial designs were analyzed with F1-LD-F1 models, independent one-way comparisons with Kruskal–Wallis tests, and dependent one-way comparisons with Friedman tests. For the F1-LD-F1 models, week main effects and cohort/condition-week interactions are reported as F-approximated ANOVA-Type Statistic (ATS) values and main effects of cohort/condition as ATS values with box-corrected degrees of freedom. Relative treatment effects (RTEs) are reported for all significant main effects and interactions (Table 3). Holm-Bonferroni corrections were applied for all pairs of post hoc comparisons. Only post hoc tests relevant to main findings are described in figures. Confidence intervals (95%) were estimated using percentile bootstrapping (2000 resamples) on the Hodges-Lehmann (HL) location shift estimate or using the Student’s t-distribution where more appropriate. Repeated measures correlations were performed using ‘rmcorr’ in R (Bakdash & Marusich, 2024) which fits subject-specific regression lines using a common slope and individual intercepts. This approach estimates the relationship between two variables across repeated observations while accounting for individual differences by fitting subject-specific intercepts while estimating a common slope across all subjects. For figures, this common slope through the grand mean is shown. All data are presented and plotted as mean ± standard error of the mean. Significance in figures is denoted as ns for not significant; * p< 0.05; ** p < 0.01; *** p < 0.001.

## Author Contributions

B.E.M performed research. B.E.M and G.O.S designed research, analyzed data, and wrote the paper. G.O.S secured funding.

## Acknowledgments

This work was supported by funds to G.O.S from The Charles E. Kaufman Foundation of The Pittsburgh Foundation (KA2024-144010) and the National Institutes of Health (AA028579, GM162859).We thank Jake Gronemeyer for help with the virtual reality tasks, Yuting Bai and Emily Le for help with animal work, and Dr. Cherish Ardinger, Dr. Kyle Jenks, and Dr. Peter Rupprecht for valuable feedback on the manuscript.

## Conflicts of Interest

The authors declare no conflicts of interest.

## Data Availability Statement

All data and code generated in this study are available from the corresponding author upon reasonable request.

## Notes

### Competing Interest Statement

The authors have declared no competing interest.

